# Bispecific antibody-drug conjugates targeting EGFR and LGR5 exert potent antitumor activity in colorectal cancer models

**DOI:** 10.64898/2026.06.22.733843

**Authors:** Peyton C. High, Maya G. Cappellino, Sean P. Sullivan, Tiffani A. Blackburn, Cara Guernsey-Biddle, Zhengdong Liang, Kendra S. Carmon

## Abstract

Colorectal cancer (CRC) remains a significant contributor to cancer-associated deaths worldwide, indicating the need for new therapeutic targets and modalities. Antibody-drug conjugates (ADCs) have demonstrated remarkable potential for the treatment of various cancer types, although their efficacy as monotherapies is often limited by insufficient targeting of tumor heterogeneity, dose-limiting toxicities, and drug resistance. Accordingly, multi-targeting therapeutic strategies, such as bispecific ADCs (bsADCs), which simultaneously target two cancer-associated antigens or non-overlapping epitopes on the same antigen, may prove more effective at overcoming resistance and eliminating tumors compared to monospecific ADCs. In this work, we describe the development of EGFR:LGR5 bispecific antibodies (bsAbs) and bsADCs. EGFR:LGR5 bsAbs were shown to internalize to the lysosome to a greater extent than EGFR- and LGR5-targeting monoclonal antibodies (mAbs) and drive EGFR lysosomal degradation in an LGR5-mediated fashion. However, EGFR:LGR5 bsAbs exerted suboptimal cytotoxicity in CRC cell lines. We therefore engineered an EGFR:LGR5 bsADC that demonstrated 100- to 1000-fold enhanced efficacy over a previously developed LGR5-targeting monospecific ADC (8E11-CPT2) with an identical linker-payload in CRC cell lines of various genetic backgrounds and EGFR and LGR5 expression levels. EGFR:LGR5 bsADC potency was strongly correlated with cell line sensitivity to the CPT2 payload. EGFR:LGR5 bsADC induced tumor regression in select *RAS*^MUT^ CRC xenograft models and demonstrated superior antitumor activity and prolonged survival benefit in all evaluated models versus EGFR mAb cetuximab (CTX), bsAb, and 8E11-CPT2. These findings strongly support the further development of EGFR and LGR5 dual-targeting approaches for CRC and other EGFR- and LGR5-expressing malignancies.

**One Sentence Summary:** EGFR:LGR5 bsADCs exert robust antitumor activity and outperform EGFR:LGR5 bsAb and LGR5 monospecific ADC in *RAS*^WT^ and *RAS*^MUT^ colorectal cancer models.

## INTRODUCTION

Antibody-drug conjugates (ADCs) represent a highly promising class of anti-cancer therapeutics, with 15 currently approved by the FDA and hundreds more in clinical development (*1*). ADCs such as HER2-targeting trastuzumab deruxtecan (T-Dxd) have revolutionized treatment standards for breast cancer (*2–5*), lung cancer (*6, 7*), and gastric cancer (*8–10*), in both frontline and treatment-refractory settings, exemplifying the vast potential of this therapeutic class for various tumor types. T-Dxd also showed promising anti-tumor activity in HER2-amplified metastatic colorectal cancer (mCRC), including those with *RAS* and/or *PIK3CA* mutations (*11*), suggesting that topoisomerase 1 inhibitor-conjugated ADCs are a promising option to improve mCRC outcomes irrespective of mutations that commonly confer resistance to current standards of care. However, only 2-4% of mCRC patient tumors overexpress HER2 (*12*), highlighting the need for improved target selection to increase the number of patients eligible to benefit from ADC therapy. Furthermore, ADC efficacy as monotherapy, particularly in solid tumors, is limited by resistance and dose-limiting toxicities (*13–18*). For this reason, ADCs are increasingly being evaluated in combination with other agents (e.g., chemotherapy, immunotherapy, targeted small-molecule and antibody-based therapies) in pre-clinical and clinical settings (*19*).

Bispecific ADCs (bsADCs), which simultaneously target two cell-surface receptors or non-overlapping epitopes on the same antigen, have also emerged as a promising option to combat resistance mechanisms encountered by traditional monospecific ADCs. In particular, several bsADCs targeting the epidermal growth factor receptor (EGFR) and other cancer-associated antigens have shown potential for the treatment of solid tumors (*20–22*). Izalontamab brengitecan (Iza-bren), for example, is an EGFR- and HER3-targeting bsADC with a topoisomerase I inhibitor payload (Ed-04; drug-to-antibody ratio (DAR) = 8) that demonstrated superior antitumor efficacy over EGFR and HER3 monospecific ADCs in pre-clinical CRC xenograft models (*23*). Iza-bren has shown promising efficacy and manageable safety profiles in late-stage clinical trials for several solid tumors and is currently in Phase 2 trials for locally advanced and mCRC in combination with PD-1-targeting mAb (NCT06008054) (*24, 25*). These findings support the development of EGFR-derived bsADCs for various tumor types.

Our group and others previously reported the development of ADCs targeting leucine-rich repeat-containing G protein-coupled receptor 5 (LGR5) (*26–31*), a marker of normal adult intestinal stem cells (ISCs) and CRC cancer stem-like cells (CSCs) with roles in supporting CRC tumorigenesis, primary tumor growth, metastatic outgrowth, and therapy resistance (*32–37*). Importantly, LGR5 is highly overexpressed in a significant majority of CRC tumors with comparably lower expression in normal tissues (*26, 28, 38*). Furthermore, genetic or radiation-induced ablation of Lgr5-expressing ISCs in adult mice had no effect on intestinal morphology nor survival, suggesting that therapeutic targeting of LGR5 is likely safe (*39–41*). LGR5-targeting ADCs incorporating microtubule or topoisomerase I inhibitors were well-tolerated and exerted significant anti-tumor efficacy in CRC xenograft models (*26–28*). However, tumors eventually relapsed, likely due to contributions from LGR5^-^ cell populations that evade ADC treatment and fuel tumor regrowth (*26, 27, 32, 35*). We showed EGFR-targeting therapies such as cetuximab (CTX), which is approved for *KRAS*^WT^ metastatic CRC, increased LGR5 protein expression in CRC cell lines, tumor organoids, and patient-derived xenograft (PDX) models (*27*). Combination of CTX with LGR5-targeting ADCs significantly reduced tumor burden and extended survival as compared to LGR5-targeting ADC and CTX monotherapies in *RAS*^MUT^ models (*27*). Consistently, recent reports have demonstrated that oncogenic Kras and Braf inhibition increased *Lgr5* levels to drive therapeutic resistance in CRC mouse models and that simultaneous restriction of *Lgr5* acquisition significantly enhanced antitumor efficacy of Kras or Braf inhibitors (*42, 43*). These findings are in line with an accumulating body of evidence that elevated Wnt and mitogen-activated protein kinase (MAPK) signaling, respectively, define functionally distinct cell populations in CRC and that cell state shifts between these populations can mediate therapeutic resistance (*42–46*). Co-targeting of Wnt and MAPK pathways via LGR5 and EGFR may therefore serve as a more effective therapeutic approach as compared to traditional mono-targeting approaches.

Consistent with previous studies demonstrating the therapeutic potential of dual-targeting EGFR and LGR5, an EGFR:LGR5 bsAb, petosemtamab (MCLA-158), is in late-stage clinical development and shows promise for treating EGFR- and LGR5-expressing tumors, including mCRC (NCT03526835). Preclinical studies showed that petosemtamab internalizes and degrades EGFR in an LGR5-dependent fashion (*47*). Importantly, petosemtamab significantly reduced tumor growth and metastasis as compared to CTX in *KRAS*^MUT^ CRC patient-derived orthotopic xenografts (*47*). Petosemtamab demonstrated minimal toxicity against LGR5-expressing ISCs and significantly less cytotoxicity in normal mucosa-derived organoids as compared to tumor organoids whereas CTX had identical cytotoxicity in both normal mucosa and tumor organoid models (*47*). The enhanced tumor selectivity of petosemtamab is likely due to the differential overexpression of LGR5 in CRC, given that EGFR is highly expressed in both normal tissue and tumors. Taken together, petosemtamab may confer both enhanced efficacy and tolerability as compared to EGFR-targeting mAbs.

Here, we describe the development of two EGFR:LGR5 bsAbs (E⨯L-1 and E⨯L-2) that internalize to a greater extent than EGFR- and LGR5-targeting mAbs and promote EGFR lysosomal degradation in an LGR5-driven manner. EGFR:LGR5 bsAbs exhibited minimal cytotoxicity in CRC cell lines, warranting the development of EGFR:LGR5 bsADCs. We generated a camptothecin derivative (CPT2)-conjugated EGFR:LGR5 bsADC (E⨯L-1-CPT2) via a site-specific chemoenzymatic approach (*27, 48, 49*). E⨯L-1 was selected over E⨯L-2 for bsADC generation and in vivo studies given that E⨯L-1 is structurally similar to clinical-stage petosemtamab and incorporated a common light chain versus two distinct light chains in E⨯L-2, thereby reducing potential for erroneous light chain mispairing. E⨯L-1-CPT2 demonstrated high specificity and 100- to 1000-fold enhanced potency compared to 8E11-CPT2, a previously-described LGR5 ADC with identical-linker payload (*27*), in EGFR- and LGR5-expressing cancer cell lines. E⨯L-1-CPT2 exerted potent efficacy in *RAS*^MUT^ CRC xenograft models with significantly enhanced anti-tumor activity and survival benefit versus CTX, E⨯L-1 bsAb, and 8E11-CPT2. Importantly, in select xenograft models, E⨯L-1-CPT2 treatment induced substantial tumor regression. These findings highlight dual-targeting of EGFR and LGR5 as a highly effective option for the treatment of CRC and other EGFR- and LGR5-expressing malignancies.

## RESULTS

### Analysis of EGFR and LGR5 expression in a panel of CRC models

We first verified expression of EGFR and LGR5 in a panel of gastrointestinal cancer cell lines and CRC patient-derived xenograft (PDX) models by Western blot and analysis of Cancer Cell Line Encyclopedia RNA-seq dataset (**Fig. 1A, S1A**) (*50*). Notably, EGFR was highly expressed in all models except for SW620 cells. SW1116, HT-29, and HCT116 cells express low to undetectable LGR5. While EGFR and LGR5 were both highly expressed across the majority of the cancer cell lines and PDX models tested, analysis of The Cancer Genome Atlas (TCGA) colorectal adenocarcinoma patient dataset revealed that *EGFR* levels are significantly reduced in tumors as compared to matched normal tissue (**Fig. 1B**). *LGR5*, however, was significantly increased in tumors as compared to matched normal tissues (**Fig. 1B**), as has been previously reported (*26, 28*). Dual-targeting of EGFR and LGR5 may therefore improve tumor selectivity and reduce off-tumor, on-target effects associated with EGFR-targeting therapies (*51*).

**Fig. 1.**
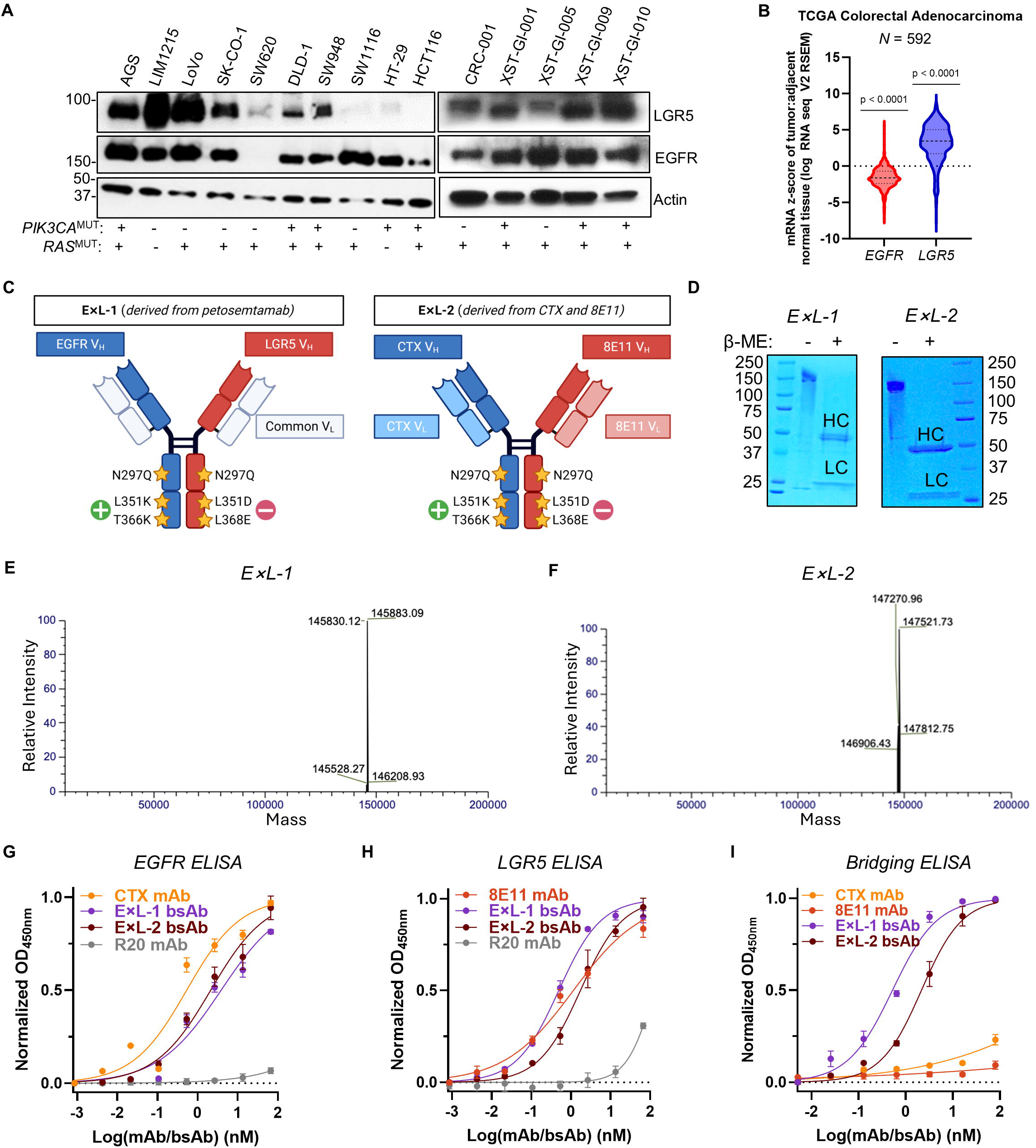
Generation of two EGFR:LGR5 bsAbs: E⨯L-1 and E⨯L-2. (**A**) LGR5 and EGFR protein expression in a panel of gastrointestinal cancer cell lines and CRC PDX models of various *PIK3CA* and *RAS* mutation statuses. (**B**) mRNA z-score of *EGFR* and *LGR5* expression in tumor versus normal adjacent tissue from the TCGA Colorectal Adenocarcinoma cohort. Significant deviation from 0 (no change in mRNA expression) was determined using Wilcoxon Signed Rank Test. (**C**) Schematic representation of E⨯L-1 and E⨯L-2 bsAbs. (**D**) Coomassie staining of SDS-PAGE for E⨯L-1 and E⨯L-2 bsAbs under native (−) and reducing (+) conditions. HC: heavy chain; LC: light chain. LC-MS/MS analysis of (**E**) E⨯L-1 and (**F**) E⨯L-2 bsAbs. (**G**) EGFR and (**H**) LGR5 single-target ELISAs performed to assess E⨯L-1 and E⨯L-2 antigen affinities as compared to CTX and 8E11 mAbs, respectively. (**I**) Dual-target bridging ELISA confirms simultaneous target engagement for E⨯L-1 and E⨯L-2 bsAbs, but not CTX or 8E11 mAbs. Antibodies were added starting at 83.3 nM and in 3-fold serial dilution for ELISAs.

### Generation and characterization of two EGFR:LGR5 bsAbs

We generated two EGFR:LGR5 bsAbs using the “DEKK” method, which involves the incorporation of negatively-charged (L351D; L368E) and positively-charged (L351K; T366K) amino acid mutations into the CH3 regions of the antibody Fc to facilitate heavy chain heterodimerization (*52*). Two EGFR:LGR5 bsAbs were generated to evaluate the reproducibility of EGFR:LGR5 bsAb-mediated effects with bsAbs integrating different structures and variable regions. E⨯L-1 bsAb was generated by subcloning the EGFR and LGR5 heavy chain variable regions (VH) from petosemtamab into human IgG1 framework incorporating L351K/T366K and L351D/L368E mutations in the EGFR and LGR5 heavy chains, respectively. E⨯L-1 also incorporates the petosemtamab common light chain. In contrast, commercially-produced petosemtamab contains L351K/T366K and L351D/L368E in the opposite heavy chains, LGR5 and EGFR, respectively, and its Fc region is engineered to enhance immune-mediated effects (e.g., antibody-dependent cellular cytotoxicity (ADCC)) by incorporating low levels of fucose (*53*). E⨯L-2 bsAb was similarly generated utilizing CTX and 8E11 mAb (*28*) constructs and incorporating L351K/T366K and L351D/L368E mutations into the CTX and 8E11 Fc regions, respectively. Notably, whereas E⨯L-1 has two distinct heavy chains and one common light chain, E⨯L-2 has two heavy chains and two light chains. Both E⨯L-1 and E⨯L-2 also incorporate an N297Q mutation in the CH2 Fc domain to facilitate site-specific drug-payload conjugation. E⨯L-1 and E⨯L-2 bind to domain III of EGFR, overlapping with the EGF ligand binding site (*47, 54*). E⨯L-1 binds to the N-CAP and first leucine-rich repeat (LRR) of LGR5 (*47*), proximal to the RSPO binding site. The exact epitope to which E⨯L-2 binds LGR5 is unknown, however, it has been shown that 8E11 mAb does not block RSPO binding to LGR5 (*28*), indicating that E⨯L-2 binds a region other than the RSPO binding site (LRRs 3-9).The structures of E⨯L-1 and E⨯L-2 are depicted in **Fig. 1C**.

E⨯L-1 and E⨯L-2 bsAbs were produced and purified as indicated by Coomassie staining (**Fig. 1D**) and LC-MS/MS (**Fig. 1E, F**). E⨯L-1 and E⨯L-2 binding to EGFR and LGR5 was confirmed by enzyme-linked immunosorbent assay (ELISA) (**Fig. 1G, H**). E⨯L-1 and E⨯L-2 had similar KD values for EGFR (E⨯L-1: 5.69 +/− 1.35 nM; E⨯L-2: 3.98 +/− 1.91 nM) and LGR5 (E⨯L-1: 1.36 +/− 0.87 nM; E⨯L-2: 2.10 +/− 0.37 nM). The KD value for LGR5 mAb 8E11 (1.29 +/− 0.14 nM) was similar, whereas the value for EGFR mAb CTX (0.39 +/− 0.25 nM) was approximately 10-fold lower, which may be attributable to bivalent EGFR binding and/or differences in the complementarity-determining regions of the antibodies. (**Table S1**). R20 mAb (Rituximab), targets CD20 and was used as an isotype, non-targeting control mAb (*27*). A dual-target bridging ELISA (*55*) was subsequently performed to confirm simultaneous receptor engagement. E⨯L-1 and E⨯L-2 showed dose-dependent increases in EGFR:LGR5 engagement whereas CTX and 8E11 mAbs did not (**Fig. 1I**), indicating that EGFR:LGR5 bsAbs simultaneously bind EGFR and LGR5.

### EGFR:LGR5 bsAbs exhibit improved internalization over mAbs

Having confirmed that E⨯L-1 and E⨯L-2 bind to EGFR and LGR5, we next examined their effect on receptor internalization via immunocytochemistry (ICC). Both E⨯L-1 and E⨯L-2 exhibited co-localization with lysosome-associated membrane protein 1 (LAMP1) in EGFR^+^/LGR5^+^ LIM1215 and DLD-1 cells following 1-hour treatment at 37°C (**Fig. 2A-B**). BsAbs showed both membrane surface binding and internalized puncta, likely attributable to EGFR and LGR5 binding, respectively, as well as LGR5-mediated EGFR:LGR5 co-internalization. This is supported by CTX displaying strong membrane localization with some internalized puncta whereas 8E11 almost exclusively shows punctate distribution (**Fig. S2A-B**) (*26, 27, 56, 57*). Still, we observed E⨯L-1 and E⨯L-2 internalization in EGFR^+^/LGR5^-^ HCT116 cells, though dramatically reduced compared to EGFR^+^/LGR5^+^ cells (**Fig. 2C**). This is consistent with previous reports of slow EGFR mAb-mediated internalization (*58*) and suggests that LGR5 drives a majority of bsAb internalization. Consistently, in EGFR^-^/LGR5^+^ SW620 cells, bsAbs exhibited predominantly LGR5-associated punctate distribution (**Fig. 2D**). To evaluate the extent of bsAb lysosomal internalization as compared to EGFR and LGR5 mAbs, we labeled CTX, 8E11, E⨯L-1, and E⨯L-2 with a pH-sensitive dye, pHrodo Red, which fluoresces upon lysosomal internalization due to increased acidification along the endolysosomal trafficking pathway. In both LIM1215 and DLD-1 cells, we observed significantly enhanced internalization for E⨯L-1 and E⨯L-2 bsAbs as compared to CTX and 8E11 mAbs following 48-hour incubation (**Fig. 2E-F**). Minimal signal was observed for 8E11 as compared to CTX, likely due to the rapid and constitutive internalization and degradation of LGR5 as compared to EGFR (*27, 57, 59*). Together, these results demonstrate that EGFR:LGR5 bsAbs internalize to the lysosome to a greater extent than EGFR- and LGR5-targeting mAbs and that this internalization is likely driven by LGR5.

**Fig. 2.**
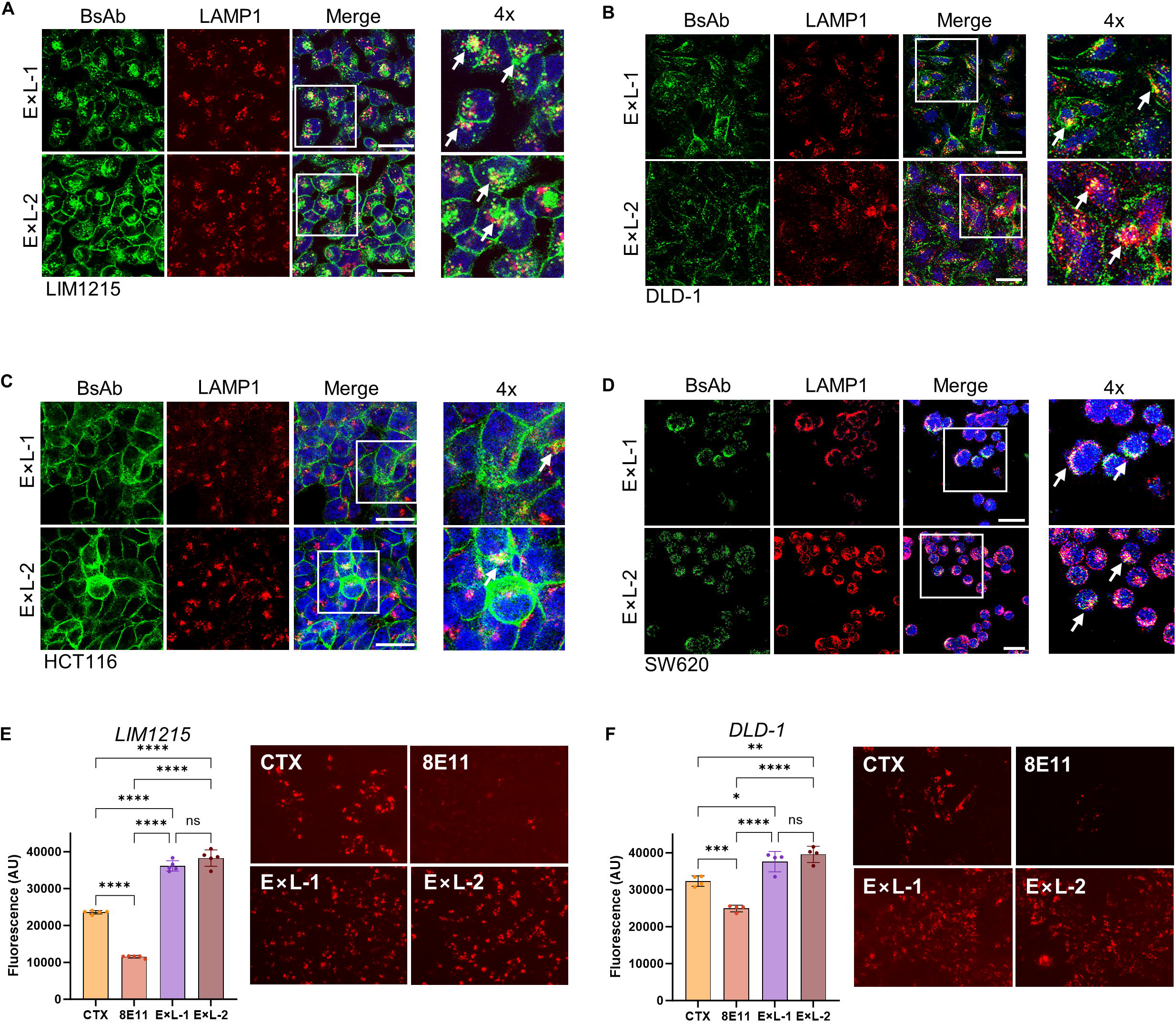
Characterization of EGFR:LGR5 bsAb internalization dynamics. ICC staining of E⨯L-1 and E⨯L-2 bsAbs following 1-hour internalization at 37°C in (**A**) EGFR^+^/LGR5^+^ LIM1215 and (**B**) DLD-1 cells, (**C**) EGFR^+^/LGR5^-^ HCT116 cells, and (**D**) EGFR^-^/LGR5^+^ SW620 cells. Arrows denote select regions of co-localization with the lysosome-specific marker LAMP1. Scale: 25 µm. (**E**) LIM1215 and (**F**) DLD-1 cells were incubated for 48 hours with 30 nM pHrodo red-labeled CTX, 8E11, E⨯L-1, or E⨯L-2. Fluorescence intensity was normalized to maximum signal-to-noise ratio and detected using Tecan Infinite M1000 plate reader. AU: arbitrary units, Significance (ANOVA): \**p*<0.05; \*\**p*<0.01; \*\*\**p*<0.001; \*\*\*\**p*<0.0001.

### EGFR:LGR5 bsAbs drive EGFR lysosomal degradation in an LGR5-dependent fashion

Having demonstrated that E⨯L-1 and E⨯L-2 internalize to the lysosome, we next examined their effect on receptor levels in EGFR- and LGR5-expressing CRC cells. Both E⨯L-1 and E⨯L-2 promoted a time-dependent reduction in EGFR levels whereas LGR5 levels were largely unchanged, if not slightly reduced (**Fig. 3A-B**). Conversely, treatment with CTX and 8E11 mAbs, individually and in combination, had no effect on EGFR levels and increased LGR5 expression (**Fig. S2C**) (*27*), indicating that EGFR downregulation requires simultaneous engagement of EGFR and LGR5. To determine the dependency of EGFR downregulation on LGR5 expression, we evaluated the effect of bsAb treatment on receptor levels in LGR5-low parental HT-29 cells and LGR5-overexpressing HT-29 cells. EGFR levels did not decrease in parental HT-29 cells following bsAb treatment (**Fig. 3C**). However, LGR5 overexpression resulted in E⨯L-1- and E⨯L-2-mediated EGFR downregulation, indicating that LGR5 enhances bsAb-mediated EGFR reduction (**Fig. 3C**). Taken together, these findings demonstrate that EGFR:LGR5 bsAbs reduce EGFR expression in an LGR5-driven manner.

**Fig. 3.**
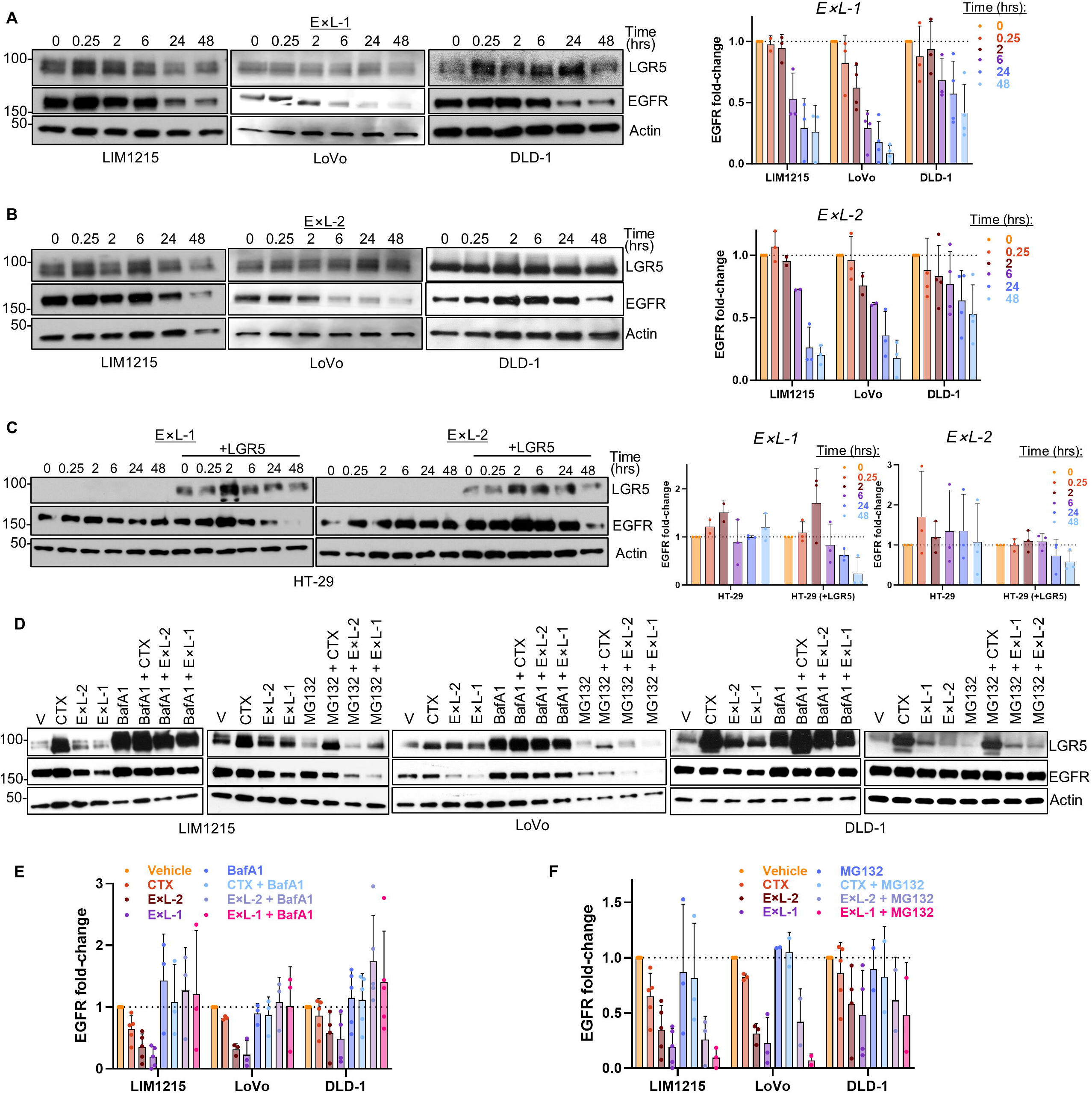
Effects of EGFR:LGR5 bsAbs on receptor expression in CRC cells. EGFR^+^/LGR5^+^ LIM1215, LoVo, and DLD-1 cells were treated with 30 nM (**A**) E⨯L-1 or (**B**) E⨯L-2 for the indicated time durations, resulting in EGFR downregulation from 6 to 48 hours. (**C**) LGR5-low or LGR5-overexpressing HT-29 cells were treated with 30 nM E⨯L-1 or E⨯L-2 for the indicated time durations. (**D**) LIM1215, LoVo, and DLD-1 cells were treated for 24 hours with 30 nM CTX, E⨯L-1 or E⨯L-2 in the presence or absence of 100 nM bafilomycin A1 (BafA1) lysosome inhibitor or 500 nM MG132 proteasome inhibitor. BafA1 or MG132 was added 15-minutes prior to antibody treatment. Quantifications of EGFR fold-change following treatment with CTX, E⨯L-1, or E⨯L-2 with and without (**E**) BafA1 or (**F**) MG132 pre-treatment

Previous works have shown LGR5 undergoes basal, constitutive internalization and trafficking to lysosomes (*27, 57, 59*). As we showed EGFR:LGR5 bsAbs traffic to lysosomes (**Fig. 2A-D**), we hypothesized that EGFR:LGR5 bsAbs drive EGFR downregulation through LGR5-mediated internalization and lysosomal degradation. To test this hypothesis, we treated LIM1215, LoVo, and DLD-1 cells with CTX, E⨯L-1 bsAb, or E⨯L-2 bsAb alone or in the presence of lysosomal inhibitor Bafilomycin A1 (BafA1; **Fig. 3D-E**). BsAb-mediated EGFR downregulation was partly to fully blocked in the presence of BafA1 while LGR5 levels were increased in the presence of BafA1 irrespective of mAb/bsAb treatment (**Fig. 3D-E**). Conversely, MG132-mediated proteasomal inhibition did not block bsAb-mediated EGFR degradation (**Fig. 3D, F**), confirming this process is lysosome-driven. Notably, E⨯L-1 treatment consistently induced more substantial EGFR degradation than E⨯L-2 (**Fig. 3**), which may be attributable in part to superior dual-target engagement of E⨯L-1 over E⨯L-2 (**Fig. 1I**). These findings demonstrate that EGFR:LGR5 bsAbs hijack LGR5 constitutive internalization and trafficking to drive EGFR lysosomal degradation.

### Development of a camptothecin-derivative (CPT2)-conjugated EGFR:LGR5 bsADC

Next, we evaluated the effects of EGFR:LGR5 bsAbs on CRC cell viability. E⨯L-1 and E⨯L-2 had moderate cytotoxic effects in EGFR- and LGR5-expressing *KRAS*^WT^ LIM1215 and *KRAS*^MUT^ LoVo and, to a lesser extent, DLD-1 cells (**Fig. 4A-C**). The cytotoxicity profiles of E⨯L-1 and E⨯L-2 were largely consistent with the effects induced by CTX mAb (**Fig. 4A-C**). These data therefore rationalize the development of EGFR:LGR5 bsADCs to enhance the potency and efficacy of EGFR:LGR5 bsAbs.

**Fig. 4.**
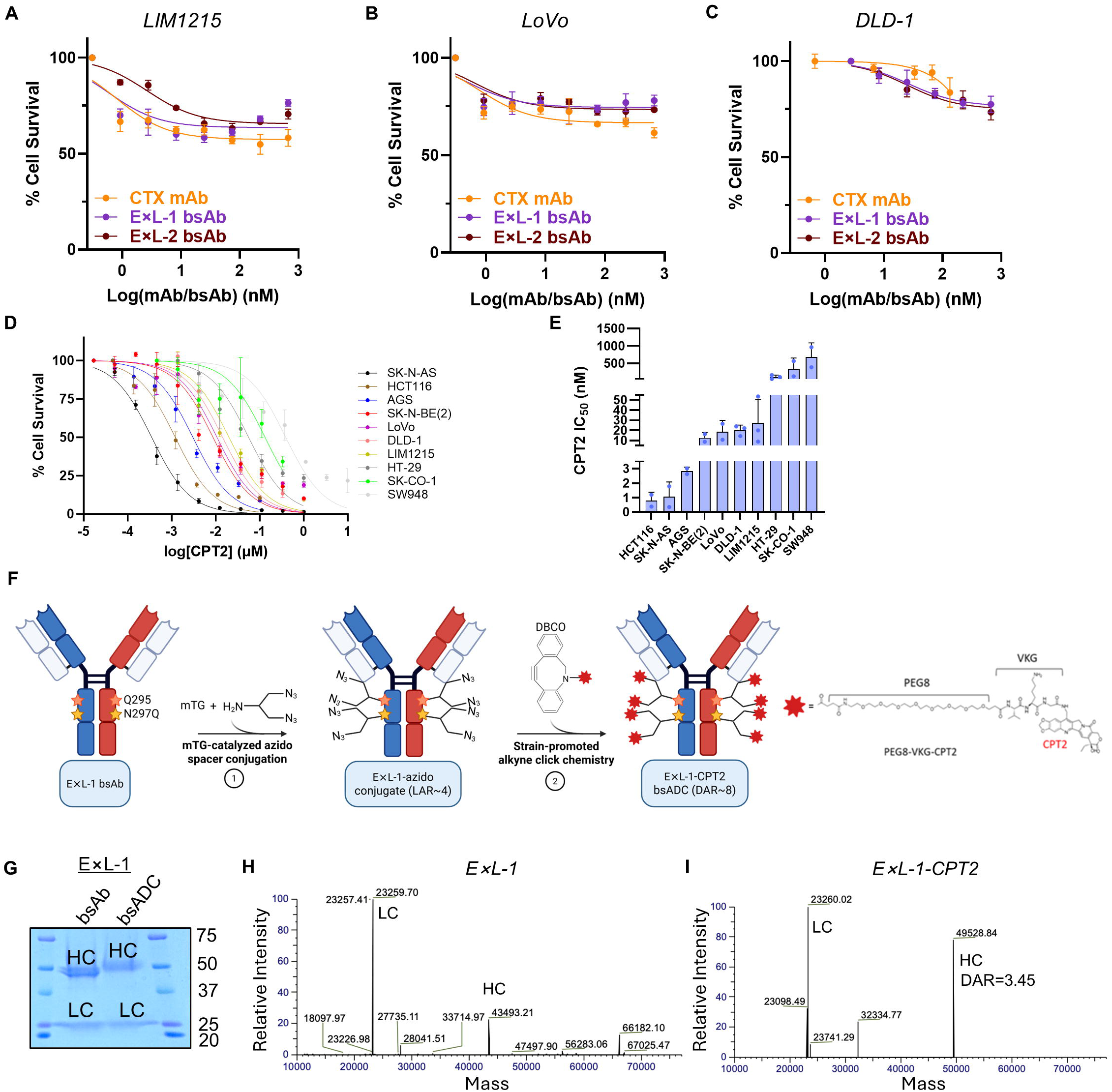
Generation of a camptothecin-conjugated EGFR:LGR5 bsADC: E⨯L-1-CPT2. (**A**) *KRAS*^WT^ LIM1215, (**B**) *KRAS*^MUT^ LoVo, and (**C**) DLD-1 cells were treated with CTX, E⨯L-1, or E⨯L-2 for 4 days starting at 667 nM in 3-fold serial dilution. (**D**) CPT2 free payload 72-hour cytotoxicity assays performed in a panel of cancer cell lines starting from 100 nM to 10 µM in 3-fold serial dilution. Cell viability was quantified using CellTiter Glo. (**E**) CPT2 IC50 values presented as mean +/− SD. (**F**) Schematic of microbial transglutaminase (mTG)-mediated conjugation of a branched linker to Q295 and N297Q of the Fc region of E⨯L-1 bsAb followed by strain promoted alkyne-azide cycloaddition of camptothecin 2 (CPT2) payload attached to a cleavable tripeptide valine-lysine-glycine (VKG) linker to generate E⨯L-1-CPT2 ADC (DAR = 8). (**G**) Coomassie staining of E⨯L-1 bsAb and E⨯L-1-CPT2 bsADC under reducing conditions shows an increase in HC molecular weight, indicating linker-payload conjugation. (**H, I**) LC-MS/MS analysis of E⨯L-1 bsAb and E⨯L-1-CPT2 bsADC under reducing conditions.

Prior to developing an EGFR:LGR5 bsADC, we analyzed cancer cell line sensitivity to the topoisomerase I inhibitor payload CPT2 (*27, 49*). Alongside CRC cell lines, we also evaluated CPT2 sensitivity in LGR5-high AGS gastric cancer (GC) and SK-N-AS and SK-N-BE(2) neuroblastoma (NB) cells (**Fig. 1A, S3A**). CPT2 exhibited nanomolar, dose-dependent potency in all cell lines evaluated, though there was substantial variation in intrinsic CPT2 sensitivity between cell lines (**Fig. 4D-E; Table S2**). We generated an EGFR:LGR5 bsADC (E⨯L-1-CPT2) utilizing site-specific, microbial transglutaminase (mTG) chemoenzymatic click chemistry conjugation as previously described (*27, 48, 60*) (**Fig. 4F**). E⨯L-1 was selected as the bsAb backbone over E⨯L-2 given that it is structurally similar to petosemtamab, which is in clinical development, and incorporates a common light chain to better prevent mispairing. Briefly, mTG catalyzes the addition of branched, azido-functionalized spacers at two glutamine (Q) residues (Q295; N297Q) in the CH2 region followed by a strain-promoted azido-alkyne click chemistry reaction resulting in the attachment of a dibenzocyclooctyne (DBCO)-functionalized linker-payload (**Fig. 4F**). E⨯L-1-CPT2 incorporates a tripeptide (VKG) protease-cleavable linker and CPT2 payload (*49*). Successful conjugation was confirmed by increased molecular weight of the ExL-1-CPT2 heavy chain compared to E⨯L-1 bsAb on Coomassie staining (**Fig. 4G**) and by LC-MS/MS (**Fig. 4H-I**; DAR = 6.91 (3.45 per HC)).

### EGFR:LGR5 bsADC outperforms LGR5 ADC and EGFR:HER3 bsADC in cancer cell lines

EGFR:LGR5 bsADC cytotoxicity was evaluated in cancer cell lines with varied EGFR and LGR5 expression and mutational statuses. E⨯L-1-CPT2 promoted time-dependent PARP cleavage in LoVo and DLD-1 cells at 24 to 72 hours, indicating apoptosis induction (**Fig. S3A**). Furthermore, similar to E⨯L-1 and E⨯L-2 bsAbs, E⨯L-1-CPT2 induced EGFR downregulation whereas 8E11-CPT2 LGR5 ADC had no effect on EGFR expression (**Fig. S3B**). We then directly compared E⨯L-1-CPT2 efficacy to 8E11-CPT2 which was generated with the identical conjugation strategy and linker-payload (*27*). E⨯L-1-CPT2 exerted 100- to 1000-fold enhanced potency over 8E11-CPT2 in EGFR- and LGR5-expressing AGS, LIM1215, LoVo, DLD-1, SW948, and SK-CO-1 cells (**Fig. 1A, 5A-F**). E⨯L-1-CPT2 also elicited dose-dependent cytotoxicity in LGR5^-^ HCT116 cells and LGR5-low HT-29 cells (**Fig. 1A, 5G-H**), indicating that E⨯L-1-CPT2 can internalize and exert cytotoxicity through EGFR independent of LGR5, consistent with ICC findings (**Fig. 2C**). E⨯L-1-CPT2 paralleled 8E11-CPT2 cytotoxicity in EGFR^-^/LGR5^+^ SW620 cells (**Fig. 5I**). EGFR overexpression conferred dramatically enhanced potency to E⨯L-1-CPT2 while that of 8E11-CPT2 was unchanged (**Fig. 1A, 5I**). Similar trends were observed in EGFR- and LGR5-high AGS GC cells and SK-N-AS and SK-N-BE(2) NB cells with E⨯L-1-CPT2 demonstrating enhanced potency over 8E11-CPT2 (**Fig. 5J-K)**, supporting the applicability of E⨯L-1-CPT2 beyond GI cancers. E⨯L-1-CPT2 and 8E11-CPT2 IC50 values are shown in **Fig. 5L** and **Table S3**. Notably, we observed a strong positive correlation between bsADC and CPT2 free payload IC50s (R^2^ = 0.7207; **Fig. 5M**), suggesting payload sensitivity may be a strong predictor of bsADC efficacy. Non-targeting ADC R20-CPT2 (*27*), generated using the same linker-payload and conjugation strategy as 8E11-CPT2 and E⨯L-1-CPT2, had no effect in CRC cell lines (**Fig. S3C**), indicating 8E11-CPT2 and E⨯L-1-CPT2 cytotoxic effects are target-dependent.

**Fig. 5.**
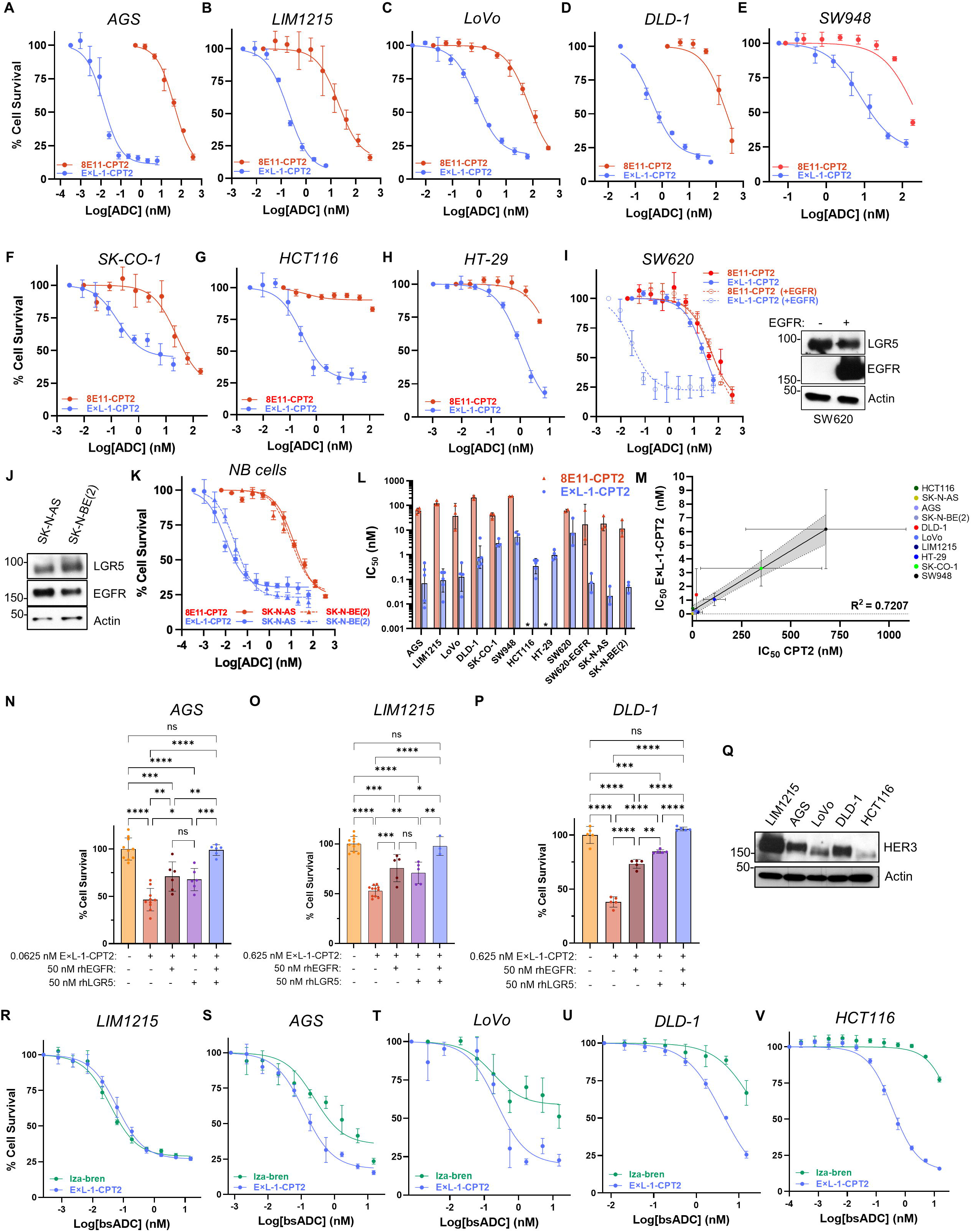
Characterization of E⨯L-1-CPT2 efficacy in cancer cell lines. In vitro cytotoxicity assays of E⨯L-1-CPT2 and 8E11-CPT2 in (**A**) AGS, (**B**) LIM1215, (**C**) LoVo, (**D**) DLD-1, (**E**) SW948, (**F**) SK-CO-1, (**G**) HCT116, (**H**) HT-29, and (**I**) SW620 +/− EGFR CRC cells. (**J**) LGR5 and EGFR expression in SK-N-AS and SK-N-BE(2) NB cells. (**K**) In vitro cytotoxicity assays of E⨯L-1-CPT2 and 8E11-CPT2 in SK-N-AS and SK-N-BE(2) cells. (**L**) Average IC50 values (+/− SD) for 8E11-CPT2 and E⨯L-1-CPT2 ADCs. (**M**) Correlation plot examining E⨯L-1-CPT2 bsADC and CPT2 free payload IC50. Correlation analyzed using simple linear regression with 95% confidence. R^2^ coefficient of determination calculated as goodness-of-fit measure. (**N**) AGS, (**O**) LIM1215, and (**P**) DLD-1 cells treated with E⨯L-1-CPT2 bsADC for four days in the presence or absence of 50 nM recombinant human (rh) EGFR and LGR5 ECDs, individually or simultaneously. Significance (ANOVA): \**p*<0.05; \*\**p*<0.01; \*\*\**p*<0.001; \*\*\*\**p*<0.0001. (**Q**) HER3 expression in LIM1215, AGS, LoVo, DLD-1, and HCT116 cancer cell lines. In vitro cytotoxicity assays comparing E⨯L-1-CPT2 and iza-bren cytotoxicity in (**R**) LIM1215, (**S**) AGS, (**T**) LoVo, (**U**), DLD-1, and (**V**) HCT116 cells. All ADC treatments were conducted for four days. Data presented as mean +/− SD. Cell viability was quantified using CellTiter Glo 2.0.

To evaluate the mechanisms of E⨯L-1-CPT2 cell-killing as determined by either EGFR or LGR5, we performed competition cytotoxicity assays wherein AGS, LIM1215, and DLD-1 cells were treated with E⨯L-1-CPT2 in the absence or presence of purified recombinant EGFR or LGR5 extracellular domains (ECDs), individually and in tandem, to block bsADC binding to either or both receptors. BsADC co-treatment with either ECD partially rescued cell survival whereas co-treatment with both EGFR and LGR5 ECDs fully rescued cell survival (**Fig. 5N-P**). These findings indicate that binding to both EGFR and LGR5 individually modulate bsADC cell-killing ability and that binding to both receptors is required for maximal efficacy.

We then directly compared E⨯L-1-CPT2 to EGFR:HER3 bsADC iza-bren, which is in Phase 2 trials for mCRC (NCT06008054) and incorporates a similar topoisomerase I inhibitor payload (Ed-04; DAR=8). E⨯L-1-CPT2 outperformed or mirrored iza-bren potency in EGFR^+^/HER3^+^/LGR5^+^ LIM1215, AGS, LoVo, and DLD-1 cells and EGFR^+^/HER3^+^/LGR5^-^ HCT116 cells (**Fig. 5Q-V**). The improved potency of E⨯L-1-CPT2 over iza-bren in LGR5^-^ HCT116 cells (**Fig. 5V**) suggests that this is not solely due to LGR5-mediated effects but may also be attributable to enhanced payload potency, superior binding and internalization through EGFR, and/or other structural differences including bsADC format, conjugation approach, or linker structure.

### EGFR:LGR5 bsADC reduces tumor burden and extends survival in CRC xenografts

E⨯L-1-CPT2 efficacy was then assessed in EGFR- and LGR5-expressing CRC cell line-derived xenograft (CDX) and PDX models. First, E⨯L-1-CPT2 was evaluated in a 5 mg/kg single-dose treatment regimen in DLD-1 CDX models. We observed significant tumor growth inhibition (TGI) of 62.9% compared to vehicle when the first vehicle-treated mouse reached maximal tumor burden (**Fig. 6A-B; S4A-B**). E⨯L-1-CPT2 treatment also extended survival as compared to vehicle treatment (**Fig. 6C**). We then evaluated 2.5 mg/kg E⨯L-1-CPT2 single- (QW) versus two-dose (QW⨯2; day 0 and day 7) regimens in an *NRAS*^MUT^ CRC-001 PDX model of rectal cancer liver metastasis (**Fig. 1A**) (*60*). Notably, both QW and QW⨯2 treatments resulted in a substantial tumor growth inhibition (TGI = 63.4% and 67.5%, respectively, when the first vehicle-treated mouse reached maximal tumor burden) as compared to vehicle (**Fig. 6D-E; S4C-D**). Though early tumor growth kinetics were not significantly different between bsADC treatment groups, we observed significantly reduced tumor volume in the QW⨯2 arm compared to QW at later time points (**Fig. 6F**). Consistently, E⨯L-1-CPT2 QW⨯2 treatment resulted in enhanced survival benefit as compared to QW treatment, both of which displayed significantly prolonged survival as compared to vehicle (**Fig. 6G**). Average body weights for DLD-1 and CRC-001 xenografts did not significantly change and are shown in **Fig. S4E-F**. Following tumor relapse, we analyzed EGFR and LGR5 expression in both CRC-001 PDX and DLD-1 CDX tumors. Contrary to the EGFR degradation induced by bsAb and bsADC treatment in vitro (**Fig. 3, S3B**), relapsed tumors expressed both EGFR and LGR5 (**Fig. S4G**), suggesting that bsADC-mediated EGFR degradation is likely transient. Importantly, these findings indicate that tumors would likely be responsive to repeated bsADC dosing, at least on the basis of retained target expression.

**Fig. 6.**
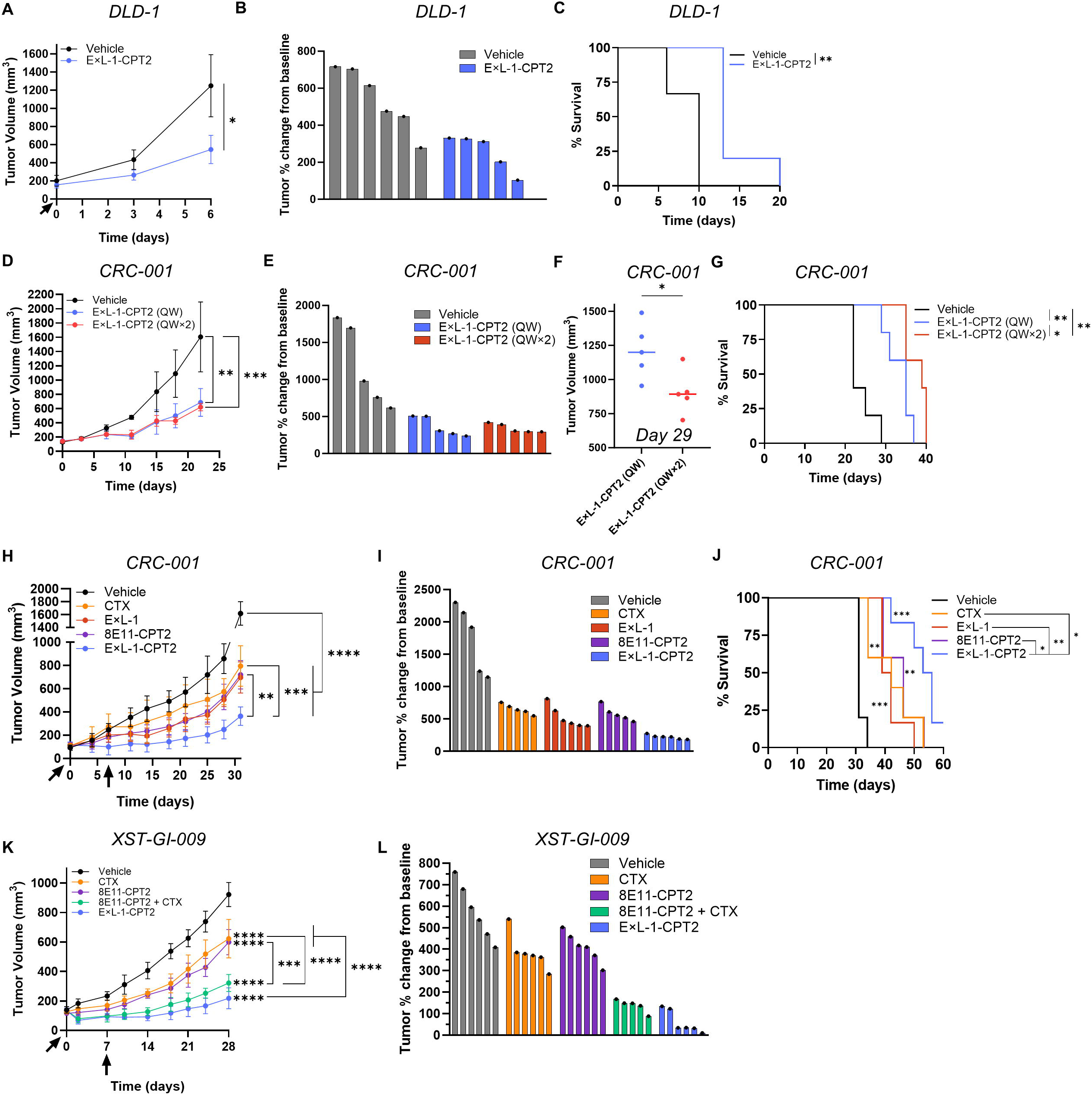
Antitumor efficacy of E⨯L-1-CPT2 in *RAS*MUT CRC xenograft models. (**A**) Tumor growth and (**B**) % change in tumor volume following E⨯L-1-CPT2 bsADC treatment in DLD-1 CDX models up to day 6 when first vehicle-treated animal reached maximal tumor burden. One treatment was administered at 5 mg/kg by IP injection (*n* = 6 for vehicle; 5 for E⨯L-1-CPT2). Statistical significance performed using ANOVA. ∗*p* < 0.05. Data presented as mean +/− SD. (**C**) Kaplan-Meier survival plot and log rank test for DLD-1 CDX study. (**D**) Tumor growth and (**E**) % change in tumor volume following QW or QW⨯2 E⨯L-1-CPT2 treatment in CRC-001 PDX models up to day 22 when first vehicle-treated animal reached maximal tumor burden. BsADC was administered at 2.5 mg/kg by IP injection (*n* = 5/group). (**F**) Tumor volume for CRC-001 bsADC-treated mice at day 29 when the first QW-treated animal reached maximal tumor burden. (**G**) Kaplan-Meier survival plot and log rank test for 2.5 mg/kg CRC-001 PDX study. (**H**) Tumor growth and (**I**) % change in tumor volume following QW⨯2 vehicle (*n* = 5), CTX (*n* = 5), E⨯L-1 (*n* = 6), 8E11-CPT2 (*n* = 5), or E⨯L-1-CPT2 (*n* = 6) treatment in CRC-001 PDX models up to day 31 when first vehicle-treated animal reached maximal tumor burden. All treatments were administered at 5 mg/kg by IP injection. (**J**) Kaplan-Meier survival plot and log rank test for 5 mg/kg CRC-001 PDX study. (**K**) Tumor growth and (**L**) % change in tumor volume following QW⨯2 vehicle (*n* = 6), CTX (*n* = 6), 8E11-CPT2 (*n* = 6), 8E11-CPT2 + CTX combination (*n* = 5), or E⨯L-1-CPT2 (*n* = 6) treatment in XST-GI-009 PDX models up to day 28. All treatments were administered at 5 mg/kg by IP injection.

We subsequently evaluated E⨯L-1-CPT2 efficacy as compared to CTX, E⨯L-1, and 8E11-CPT2 in CRC-001 PDX models dosed QW⨯2 at 5 mg/kg. CTX (TGI = 54.9%), E⨯L-1 (TGI = 61.8%), and 8E11-CPT2 (TGI = 59.9%) all significantly reduced tumor burden as compared to vehicle and to a similar extent as one another (**Fig. 6H-I; S5A-B**). Importantly, E⨯L-1-CPT2 (TGI = 83.6%) significantly enhanced antitumor activity as compared to vehicle and all other treatment groups (**Fig. 6H-I; S5A-B**). E⨯L-1-CPT2 induced initial tumor regression in 4/6 mice with tumor regrowth observed following treatment cessation at days 11-25 (**Fig. S5A**). All treatment groups extended survival as compared to vehicle, with E⨯L-1-CPT2 treatment resulting in the greatest survival benefit (**Fig. 6J**). Consistent with previous findings, no treatment groups induced adverse changes in body weight (**Fig. S5C**).

Finally, E⨯L-1-CPT2 efficacy was assessed in an EGFR- and LGR5-high *KRAS*^MUT^/*PIK3CA*^MUT^ PDX model (XST-GI-009; **Fig. 1A**) as compared to CTX, 8E11-CPT2, and 8E11-CPT2 + CTX combination dosed once weekly at 5 mg/kg for a total of two doses. E⨯L-1-CPT2 was evaluated against 8E11-CPT2 + CTX combination as an alternative EGFR and LGR5 dual-targeting approach given our previous findings that CTX significantly enhanced the antitumor efficacy of 8E11-CPT2 in *RAS*^MUT^ PDX models (*27*). Both CTX (TGI = 36.7%) and 8E11-CPT2 (TGI = 38.4%) monotherapies induced similar antitumor effects whereas 8E11-CPT2 + CTX combination treatment (TGI = 76.1%) and E⨯L-1-CPT2 (TGI = 89.4%) had the most pronounced effects (**Fig. 6K-L, S5D-E**). Substantial tumor regression was noted in all mice in both CTX + 8E11-CPT2 combination treatment (n=5; 1/5 until day 10; 1/5 until day 14; 2/5 until day 18; 1/5 until day 21) and E⨯L-1-CPT2 bsADC (n=6; 2/6 until day 18; 1/6 until day 21; 1/6 until day 24; 2/6 until day 28). Furthermore, as observed with all other xenograft studies, no treatment group induced adverse changes in body weight (**Fig. S5F**). Taken together, these results highlight the strong potential of EGFR and LGR5 dual-targeting approaches to treat CRC.

## DISCUSSION

While ADCs have greatly transformed standards of care for multiple cancer types, their clinical efficacy is limited by toxicities and resistance mechanisms such as loss of target expression. BsADCs therefore present a potentially improved therapeutic option by enhancing tumor specificity and targeting of tumor heterogeneity. In support of this, several bsADCs have already advanced to clinical trials, including HER2 biparatopic ADCs (*61–63*) as well as multiple ADCs targeting EGFR and other cancer-associated antigens such as HER3 (*21, 24, 25*), c-MET (*22, 64, 65*), and MUC1 (*66*). Among these, the EGFR:HER3 bsADC iza-bren is the furthest advanced, currently in Phase 2 trials for mCRC and Phase 3 trials for several other solid tumors (*21, 24, 25*). In addition to EGFR:LGR5 bsADCs, the EGFR:LGR5 bsAb petosemtamab is currently being investigated in *RAS*/*RAF*^WT^ mCRC in combination with FOLFOX/FOLFIRI in first- and second-line (1/2L), anti-EGFR therapy naïve settings or as a monotherapy in third-line or later (3L+) settings, including after previous anti-EGFR therapy (NCT03526835). Interim analysis from these trials has demonstrated promising clinical efficacy with manageable toxicity in 1L, 2L, and 3L+ mCRC patients (*67*). Given the clinical success of EGFR-targeting bsADCs and petosemtamab, there is strong rationale supporting the development of EGFR:LGR5 bsADCs.

Dual-targeting of EGFR and LGR5 is of particular relevance for CRC given the clinical success of EGFR-targeting therapies in *KRAS*^WT^ mCRCs (*68*) and the functional roles of LGR5 in CRC tumorigenesis (*33*), progression (*32, 35*), metastasis (*32, 69*), and therapeutic resistance (*34, 42, 43, 70*). LGR5 has also been shown to be upregulated following treatment with EGFR-, KRAS-, and BRAF-targeting therapies (*27, 35, 42–45, 47, 71, 72*) and is capable of driving therapeutic resistance and tumor relapse. *Lgr5* ablation resulted in dramatically enhanced efficacy of EGFR and KRAS-targeting agents (*35, 43*), further supporting LGR5 as a rational target in combination with MAPK pathway-targeting therapies. In addition to the enhanced efficacy conferred by dual-targeting of EGFR and LGR5, we also anticipate improved tolerability of EGFR:LGR5 bsAbs and bsADCs over EGFR-targeting approaches due to the differential overexpression of LGR5 in tumors as compared to ubiquitous EGFR expression in normal tissue and tumors (*26*). This is supported by the preclinical characterization of petosemtamab, which demonstrated improved selectivity for tumor organoids versus normal intestinal organoids compared to CTX (*47, 53*). Notably, the lack of improved efficacy of E⨯L-1 over CTX observed in this study (**Figs. 4A-C; 6H-J**) contrasts with findings from preclinical characterization of petosemtemab, which demonstrated superior antitumor effects over CTX (*47*). The divergent findings may be attributed to the models utilized in this study that do not effectively capture immune-mediated effects, which are thought to be an important mechanism of action for petosemtemab (*47, 53*). Future studies therefore ought to evaluate E⨯L-1 and E⨯L-1-CPT2 in immune-competent models as compared to CTX to fully decipher the contributions of immune-mediated cytotoxicity of petosemtamab versus CTX, E⨯L-1, E⨯L-1-CPT2. Notably, E⨯L-1 incorporates an N297Q mutation that reduces Fc effector function (*73*) and lacks other Fc effector function-enhancing mutations that are present in petosemtamab, suggesting immune-mediated effects are likely to be more pronounced with petosemtamab over E⨯L-1.

Our work demonstrates that EGFR:LGR5 bsAbs promote degradation of EGFR via hijacking of constitutive LGR5-mediated lysosomal trafficking. These results inform a generalizable mechanism of extracellular targeted protein degradation wherein bsAbs are designed to target cancer driver proteins (e.g., EGFR) and a constitutively internalizing receptor (e.g., LGR5) to induce degradation of cell surface-expressed oncoproteins. Given that combination of EGFR and LGR5 mAbs does not induce EGFR downregulation (**Fig. S2C**), our results demonstrate that this mechanism requires simultaneous bsAb-mediated target engagement. Similar results have been observed with other bispecific modalities targeting EGFR and internalizing receptors asialoglycoprotein receptor (ASGPR) (*74*) and low-density lipoprotein receptor (LDLR) (*75*). While this presents an attractive approach to degrade cell surface proteins, EGFR:LGR5 bsAbs showed suboptimal cytotoxicity (**Fig. 4A-C**). These findings indicate that alternative strategies are necessary to enhance the potency of these agents, such as the development of bsADCs. We demonstrate in this work that EGFR:LGR5 bsADCs exhibit improved cytotoxicity over bsAbs due to payload-mediated cell death and we anticipate bsADCs retain the improved tumor selectivity and tolerability conferred by the bsAb backbone.

We report that EGFR:LGR5 bsADCs are highly effective and induce initial tumor regression in EGFR- and LGR5-expressing CRC models independent of *RAS* mutations. E⨯L-1-CPT2 demonstrated superior efficacy over E⨯L-1 bsAb and 8E11-CPT2 LGR5 ADC in *RAS*^MUT^ PDX models (**Fig. 6H-L**), indicating that the payload-mediated cytotoxicity and dual-targeting capacity of E⨯L-1-CPT2 confers enhanced anti-tumor activity as compared to bsAb and monospecific ADCs. E⨯L-1-CPT2 induced similar antitumor effects as 8E11-CPT2 + CTX combination treatment, suggesting EGFR and LGR5 dual-targeting approaches broadly are an effective approach to improve upon monotargeting approaches. Still, E⨯L-1-CPT2 may theoretically offer advantages of control of dosing and more predictable tolerability given it is administered as one therapy versus two. E⨯L-1-CPT2 also demonstrated similar or improved efficacy over EGFR:HER3 bsADC iza-bren in GI cancer cells (**Fig. 5R-V**), suggesting topoisomerase I inhibitor-conjugated EGFR:LGR5 bsADCs may be a viable clinical option to build upon the success of iza-bren. While E⨯L-1-CPT2 did not fully eliminate tumors, alternative dosing regimens and magnitudes should be evaluated to prolong initial tumor regression that was observed in select PDX models. Similar to monospecific ADCs, acquired resistance mechanisms likely influence continued therapeutic response including target downregulation, antigen structural mutations that impair bsAb binding, payload-mediated resistance mechanisms, and alterations in bsADC trafficking (*20*), which warrant further investigation.

While this study strongly supports EGFR:LGR5 bsADCs as a promising treatment for CRC, there are intrinsic limitations. Firstly, the EGFR-targeting arms of E⨯L-1 and E⨯L-2 do not recognize murine EGFR. Accordingly, EGFR-mediated on-target, off-tumor toxicities cannot be examined in rodent models and should be assessed in higher model organisms (e.g., cynomolgous monkeys) prior to human testing. Importantly, our previously generated LGR5 ADC with identical linker-payload and murine cross-reactivity was well-tolerated at all doses tested, up to 20 mg/kg, suggesting LGR5-mediated on- and off-target ADC toxicities are manageable up to and likely above this dose (*27*). Given that petosemtamab demonstrates improved tumor selectivity as compared to CTX in organoid models (*47*) and has shown manageable tolerability in clinical trials (*53, 67*), we anticipate E⨯L-1-CPT2 will have a similarly favorable safety profile. Improved tolerability of EGFR:LGR5 bsAbs over EGFR mAbs may be attributable to several factors. Firstly, LGR5 overexpression in tumors versus normal tissue confers improved selectivity for tumors and, therefore, decreased internalization and EGFR degradation in normal tissue (*47*). Furthermore, EGFR:LGR5 bsAbs display monovalent EGFR binding as compared to EGFR mAbs which have bivalent binding. Thus, it would be of interest to evaluate the tolerability of monovalent versus bivalent EGFR mAbs to determine the extent to which antibody valency influences toxicity associated with traditional EGFR mAbs.

As this work was primarily focused on establishing proof-of-concept of EGFR:LGR5 bsADCs as an effective therapeutic approach for CRC, we elected to evaluate E⨯L-1-CPT2 primarily in subcutaneous CRC PDX models. Future studies should be performed to evaluate the effects of EGFR:LGR5 bsADC treatment in EGFR/RAS/MAPK treatment-resistant and orthotopic models of CRC. Given that LGR5 is required for metastatic outgrowth and that petosemtamab reduced metastasis in pre-clinical CRC models, we anticipate E⨯L-1-CPT2 to have similar anti-metastatic effects. Additionally, given that MAPK-targeting therapies have been shown to increase LGR5 expression (*27, 35, 42–45*), it is plausible that EGFR:LGR5 bsADCs demonstrate significant efficacy in treatment-refractory settings. Given that E⨯L-1-CPT2 demonstrated anti-tumor efficacy in *RAS*^MUT^ models, this work suggests that EGFR:LGR5 bsADCs are a highly promising option to treat both *RAS*^WT^ and *RAS*^MUT^ CRCs and expand the classical indications for EGFR-targeting therapies, including petosemtemab, in mCRC. Beyond CRC, this work also suggests EGFR:LGR5 bsADCs may prove effective in other EGFR:LGR5 cancers, particularly head and neck cancers where petosemtamab has shown strong promise (NCT06496178; NCT06525220) (*76, 77*). Finally, our findings support the future development of LGR5-targeting bsADCs with alternative targets to improve upon the current arsenal of ADC therapies.

## MATERIALS AND METHODS

### Study Design

The studies presented above were conducted to develop and characterize the antitumor efficacy of EGFR:LGR5 bsAbs and bsADCs in CRC cell lines and xenograft models. All in vitro experiments are representative of 3 experimental replicates unless otherwise noted. In vitro cytotoxicity experiments were performed with 3-4 sampling replicates. In vivo rodent studies were performed in *nu*/*nu* and NSG mice to assess therapeutic efficacy in patient-representative models. Animal studies were carried out with IACUC approvals (AWC-23-0106 and AWC-24-0128). The studies were conducted in accordance with recognized ethical guidelines (i.e., Declaration of Helsinki, Belmont Report, and U.S. Common Rule) and appropriate approvals were obtained by the UTHealth Houston Institutional Review Board (HSC-MS-20-0327, HSC-MS-21-0074). No blinding to group allocation was performed and sample sizes were determined based on prior experience with the models to reach statistical significance. Mice were euthanized when tumor diameter reached 15 mm. All data points were included in the calculations described in these studies with no outliers excluded.

### Statistical Analysis

All data were analyzed using GraphPad Prism 10 software. Data presented as mean ± SD. In vitro viability and ELISA data were analyzed using the logistic nonlinear regression model. Statistical significance of matched *EGFR* and *LGR5* expression was analyzed with Wilcoxon signed-rank test. Significance for in vitro and in vivo experiments was analyzed using two-sided student T-test, ANOVA, and Tukey’s multiple comparison test. Kaplan-Meier analysis was performed to determine significance of treatments on survival. *p* < 0.05 was deemed significant.

### Cell lines

DLD-1, LoVo, HCT116, SW620, AGS, HT-29, SK-CO-1, SW948, SW1116, SK-N-AS, and SK-N-BE(2) cells were purchased from ATCC. LIM1215 cells were purchased from Millipore Sigma. Cell lines were authenticated utilizing short tandem repeat profiling and tested for mycoplasma. All cancer cell lines except for SK-CO-1 cells were cultured in RPMI medium (Gibco) supplemented with 10% fetal bovine serum (Gibco) and penicillin/streptomycin (Gibco). LIM1215 cells were additionally supplemented with 1 μg/mL hydrocortisone (Sigma-Aldrich), 10 μM 1-Thioglycerol (Sigma-Aldrich), 0.6 μg/mL insulin (Gibco) and 25 mM HEPES (Gibco). SK-CO-1 cells were cultured in EMEM medium (Gibco) supplemented with 10% fetal bovine serum and penicillin/streptomycin. Stable EGFR-overexpressing SW620 cells (SW620-EGFR) were generated using pCDNA6A-EGFR WT plasmid (gift from Mien-Chie Hung; Addgene #42665) (*78*). Stable LGR5-overexpressing HT-29 cells (HT-29-LGR5) were generated via lentiviral transduction of pCDH-CMV-LGR5-EF1α-Neo (gift from Qingyun Liu at UTHealth Houston). SW620-EGFR cells were maintained in 1 µg/ml puromycin (Gibco) and HT-29-LGR5cells were cultured in 400 µg/ml G418 (Gibco). All cell lines were cultured at 37°C with 95% humidity and 5% CO2.

### Plasmids and cloning

LGR5 mAb 8E11, EGFR mAb CTX and R20 mAb were generated as previously described (*27*). E⨯L-1 was generated using publicly available patented sequences from petosemtamab and E⨯L-2 was generated using CTX and 8E11 mAb sequences. DNA sequences encoding the variable heavy and light chain regions were codon optimized and synthesized (Epoch Life Science, Inc.) and subcloned into pCEP4 vector containing either human IgG1 or kappa constant regions using In-Fusion Snap Assembly (Takara). 8E11, R20, E⨯L-1, and E⨯L-2 incorporated an N297Q mutation in the IgG1 CH2 heavy chain Fc region for site-specific conjugation. E⨯L-1 and E⨯L-2 also incorporated L351K/T366K and L351D/L368E mutations via site-directed mutagenesis in the EGFR and LGR5 mAb monomer CH3 heavy chain Fc regions, respectively, to promote heavy chain heterodimerization.

### Western blot

Protein lysate was prepared using RIPA lysis buffer (ThermoFisher) supplemented with Halt protease and phosphatase inhibitors (ThermoFisher). PDX tissues were homogenized and freeze-thawed. Lysates were sonicated then centrifuged 10 min, 4°C at 13,000xg. Protein concentrations were quantified by BCA protein assay (Thermo Scientific) and lysates were diluted in reducing SDS Laemmli buffer (Thermo Scientific) and incubated at 37°C for 1 h prior to SDS-PAGE. Anti-mouse IgG and anti-rabbit IgG HRP-labeled secondary antibodies from Cell Signaling Technology were utilized with the standard ECL protocol (Cytiva). Primary antibodies used in this study include: anti-EGFR (Cat# 54359, 1:1000); PARP (Cat# 9542); and anti-β-actin (Cat# 3700, 1:5000) from Cell Signaling Technology and anti-LGR5 (Cat# ab75732, 1:1000) from Abcam. Chemical inhibitors used include BafA1 (MedChemExpress) and MG132 (MedChemExpress). Western Blot quantification was performed with Fiji ImageJ by normalizing protein band intensity to the explicated reference protein (e.g., actin). Relative protein-fold changes were calculated by dividing this value for each treatment condition by that of vehicle control.

### Immunocytochemistry

Cells were seeded in a poly-D-lysine-coated 8-well chamber slide (Corning Cat# 354632) and treated the following day. For lysosome co-localization experiments, cells were treated with 30 nM EGFR mAb (CTX), LGR5 mAb (8E11), E⨯L-1 bsAb, or E⨯L-2 bsAb at 37°C for 1 h, washed, fixed in 4% paraformaldehyde (ThermoFisher Scientific) and permeabilized in 0.1% saponin (Sigma-Aldrich). Cells were incubated with anti-LAMP1 (Cell Signaling Technology Cat# 9091, 1:400) at room temperature for 1 h, followed by anti-human-Alexa-488 (ThermoFisher, 1:200) and goat anti-rabbit-Alexa-555 (ThermoFisher, 1:200) at room temperature for 1 h. Nuclei were counterstained with TO-PRO3 (ThermoFisher, 1:1000) for 15 min. Images were acquired using confocal microscopy (Leica TCS SP5) and LAS AF Lite software.

### pHrodo Red antibody labeling

CTX, 8E11, E⨯L-1 and E⨯L-2 were conjugated to pHrodo Red dye (excitation 560 nm/emission 585 nm) using pHrodo Red antibody labeling kit (Invitrogen). Briefly, ∼100 µg of antibody was incubated with sodium bicarbonate (100mM final concentration) and pHrodo Red ester amine-reactive dye at a molar ratio of 10:1 dye:Ab for 15 minutes at room temperature. pHrodo Red ester amine-reactive dye covalently binds free lysines present on the antibody. Dye-labeled antibody was subsequently purified through gel resin at 1000xg for 5 minutes and concentration was determined by NanoDrop (ThermoFisher). Fluorescently labeled antibodies were stored at 4°C. For internalization experiments, LIM1215 and DLD-1 cells were incubated with 30 nM of each labeled antibody for 48 hours and fluorescence intensity was measured using the Tecan Infitite M1000 plate reader (Excitation wavelength: 560 nm; emission wavelength: 585 nm; excitation/emission wavelength: 10 nm).

### Antibody production

MAb and bsAb production was performed by transient expression of corresponding light and heavy chain constructs in Expi293F cells (ThermoFisher) using Polyethylenimine HCL Max reagent (Polysciences Cat# 247651). Media was collected 7 days post-transfection and mAbs were purified using protein A (Genscript Cat# L00210) affinity chromatography and eluted under acidic conditions (pH 3 glycine) into pH 8.5 TRIS. Buffer exchange to PBS was performed by centrifugation at 4000 rpm. Mabs and bsAbs were analyzed for purity and homogeneity by SDS-PAGE and LC-MS/MS and concentration was determined by NanoDrop (ThermoFisher).

### ELISAs

ELISA plates were coated with 23 nM recombinant human LGR5 ECD-Fc fusion (R&D Systems #8078-GP; LGR5 and bridging ELISA) or 29 nM recombinant human His-tagged, biotinylated EGFR ECD (R&D Systems #BT11302; EGFR ELISA) in PBS and incubated overnight at 4°C. The following day, plates were washed with PBS and blocked in 3% BSA for 1 hour at 37°C. E⨯L-1 bsAb, E⨯L-2 bsAb, CTX mAb, 8E11 mAb, or R20 mAb were then added in triplicate starting at 66.67 to 80 nM, in 5-fold serial dilutions, in 3% BSA and incubated for 1 hour at 37°C. For LGR5 ELISA, E⨯L-1 bsAb, E⨯L-2 bsAb, 8E11 mAb, and R20 mAb were biotinylated via the EZ-Link^TM^ Sulfo-NHS-LC-Biotin labeling kit (ThermoFisher) according to manufacturer instructions prior to incubation. For bridging ELISA, plates were then incubated with 29 nM biotinylated, recombinant EGFR ECD for 1 hour at 37°C. Next, plates were incubated with anti-human IgG-HRP (1:4000 in 3% BSA) for EGFR ELISA or streptavidin-HRP (1:200 in 3% BSA) for LGR5 or bridging ELISA for 1 hour at 37°C. Finally, TMB substrate was added to the plates and incubated at room temperature for 10-15 minutes or until sufficient color development. This reaction was stopped by adding 2M H2SO4 and absorbance was measured at λ = 450nM using Tecan Infinite M1000 plate reader. Plates were washed between each step with 0.1% PBST.

### Mono- and bispecific ADC production

For site-specific conjugation to Q295 and Q297, 8E11 mAb, R20 mAb, and E⨯L-1 bsAb were conjugated as previously described (*27*). Briefly, mAbs and bsAbs were incubated with diazido branched linker, N-(Amino-PEG2)-N-bis(PEG3-azide) (BroadPharm) (160 equivalents) and Activa TI Transglutaminase (final concentration 8%, Ajinomoto, Modernist Pantry) at room temperature for 16 hours. Conjugated mAb was purified by protein A column chromatography to afford a mAb-linker conjugate. Subsequently, DBCO-PEG8-VKG-CPT2 was synthesized (MedChemExpress) and added (1.5 equivalent per azide) to a solution of 8E11-linker, R20-linker, or E⨯L-1-linker conjugate and incubated 4 hours at RT. Crude products were purified by FPLC (Cytiva AKTA pure) to afford the ADCs.

### Intact protein mass analysis and spectral deconvolution

E⨯L-1 and E⨯L-2 bsAbs and E⨯L-1-CPT2 bsADC were analyzed at the Rice University Mass Spectrometry Proteomics Core Facility. Liquid chromatography tandem mass spectrometry experiments were performed on a Thermo Scientific Orbitrap Ascend equipped with a Thermo Scientific Vanquish Neo chromatography system. Protein samples were injected onto a Waters NanoEase m/z BEH C4 column (300 A pore size, 1.7 um particle size, 150um I.D. × 100mm). Proteins were eluted over a gradient of 3-65% acetonitrile over 7 minutes at a constant flow rate of 2µL/min. Spectra were acquired from m/z 500-8000 in positive ion mode at 2000V with a source fragmentation of 120V and a resolving power of 75000. Data were processed in Thermo Scientific Biopharma Finder 5.1. Spectra were averaged using a sliding window method (target spectrum width: 0.2 min; offset 25%) and deconvoluted using the ReSpect deconvolution algorithm with the following parameters: m/z range 500-8000; Peak model: intact protein; Model mass range: 10kDa-160 kDa; Charge state range: 10-100; Minimum adjacent charges: 6; Relative abundance threshold 10%. DAR values were previously reported for 8E11-CPT2 (7.94) and R20-CPT2 (8.0). DARE⨯L-1-CPT2 = 6.91.

### In vitro cytotoxicity assays

Cancer cells were plated at approximately 1,500 cells/well in 96 half-well plates (Corning). Serial dilutions of E⨯L-1-CPT2 or 8E11-CPT2 were added and incubated at 37°C for 4 days. Cell viability was measured using the CellTiter-Glo 2.0 Assay (Promega) according to the manufacturer’s protocol. Luminescence was measured using Tecan Infinite M1000 plate reader. For competition cytotoxicity assays, cells were incubated in the presence of 50 nM recombinant human EGFR and/or LGR5 ECDs (R&D) for the duration of bsADC treatment.

### In vivo efficacy studies

For bsADC efficacy studies in DLD-1 CDX models, female 6-8-week-old nu/nu (The Jackson Laboratory) mice were implanted subcutaneously with approximately 2.5 × 10^6^ cells. For CRC-001 and XST-GI-009 PDX ADC efficacy studies, female 6-8-week-old NSG (The Jackson Laboratory) mice were used. CRC-001 was from Jackson Laboratory (Cat# TM00849). Mice were implanted subcutaneously with 2–3 mm PDX fragments into the right flank. When tumors reached 100–200 mm^3^, mice were randomized based on equivalent average tumor volume for each treatment group. Mice were intraperitoneally dosed weekly once or twice at 2.5 mg/kg (CRC-001) or 5 mg/kg (DLD-1, CRC-001, XST-GI-009) bsADC as indicated. CRC-001 xenografts also included arms for CTX, E⨯L-1, and 8E11-CPT2 dosed intraperitoneally at 5 mg/kg once weekly for two total doses. XST-GI-009 xenografts included arms for CTX, 8E11-CPT2, and CTX + 8E11-CPT2 dosed intraperitoneally at 5 mg/kg once weekly for two total doses. Mice were routinely monitored for morbidity and mortality. Tumor volumes were measured bi-weekly and estimated by the formula: tumor volume = (length/2 × width^2^). Percentage of tumor growth inhibition (% TGI) was calculated using the formula [1 − (change of tumor volume in treatment group/change of tumor volume in vehicle group)] × 100 (%). Tumor % change from baseline was calculated using the formula (change of tumor volume/starting tumor volume) × 100 (%).

## Acknowledgments

We would like to thank Dr. Katelyn Baumer from the Rice University Shared Equipment Authority Mass Spectrometer Facility for assistance with running and processing LC-MS/MS analyses; Dr. Julie Rowe, Betty Arceneaux, and Dr. Karan Saluja with assistance in patient sample collection and processing; and Martha Thompson for assistance with regulatory approvals for research involving human subjects. Schematics were created using Biorender.com.

## Author contributions

K.S.C. and P.C.H. conceptualized the study. KSC and P.C.H. performed data curation. KSC and P.C.H. conducted formal analysis. K.S.C, P.C.H., M.G.C., S.P.S., T.A.B., C.G-B., and ZL conducted the investigation and generated the data. KSC and P.C.H. provided visualization of the figures. K.S.C. provided project administration and supervised the study. K.S.C. and P.C.H. provided funding support. P.C.H. wrote original manuscript. K.S.C., P.C.H., M.G.C., S.P.S., T.A.B., C.G-B. edited the manuscript. All authors read and approved the final manuscript.

## Data and materials availability

All raw data generated in this are available upon request from the corresponding authors. Publicly available data can be found from TCGA and CCLE (www.cbioportal.org).

