## Supplementary Figures S1-S5 and Tables S1-S3 for "Bispecific antibody-drug conjugates targeting EGFR and LGR5 exert potent antitumor activity in colorectal cancer models"

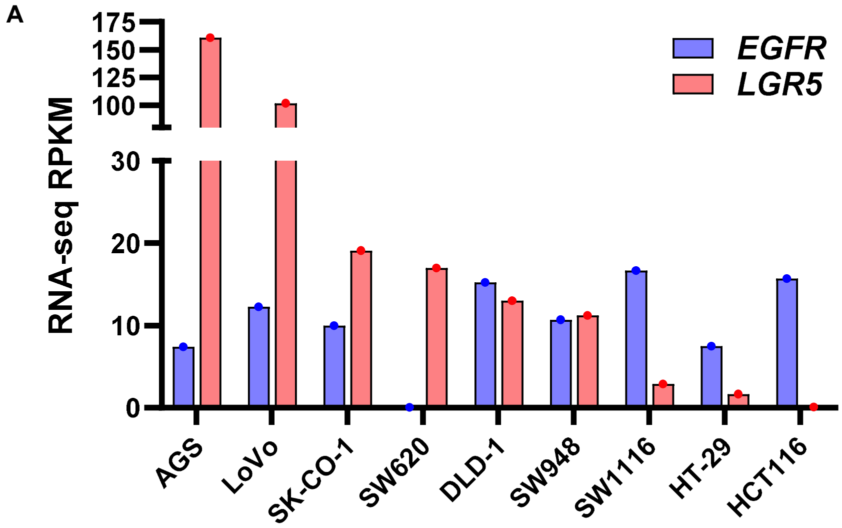


**Figure. S1. EGFR and LGR5 mRNA expression in cancer cell lines.** (**A**) mRNA expression (reads per kilobase of transcript per million mapped reads; RPKM) across gastrointestinal cancer cell lines utilized in this study. mRNA values were obtained from CCLE RNA sequencing data.

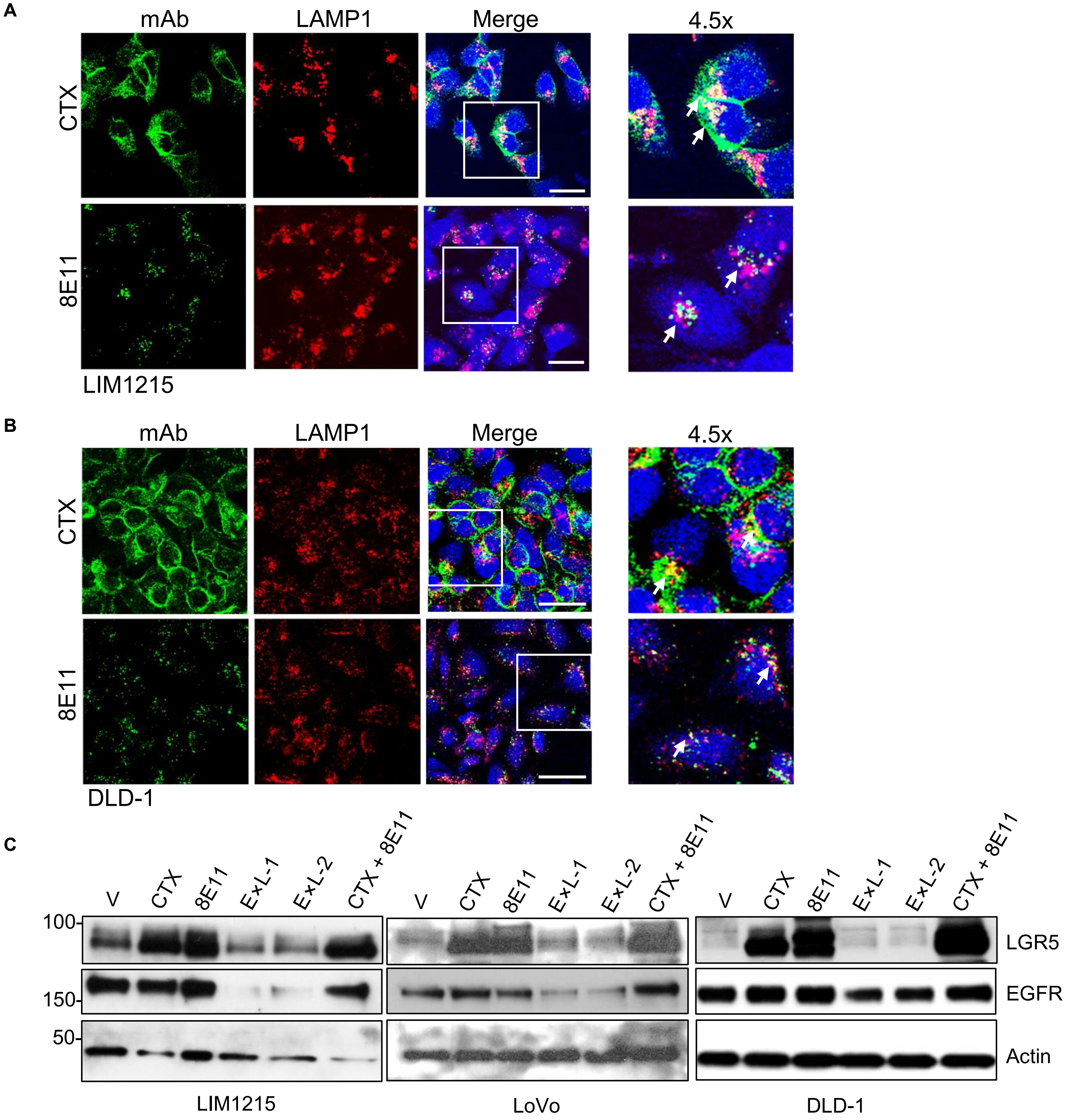


**Figure S2. CTX and 8E11 internalization and effects on receptor expression as compared to EGFR:LGR5 bsAbs.** ICC staining of CTX and 8E11 mAbs following 1-hour internalization at 37˚C in (**A**) $EGFR^{+}/LGR5^{+}$ LIM1215 and (**B**) DLD-1 cells. Arrows denote regions of co-localization with the lysosome-specific marker LAMP1. Scale: 25 µM. (**C**) LIM1215, LoVo, and DLD-1 cells were treated with 30 nM CTX, 8E11, E×L-1, E×L-2, or CTX + 8E11 combination for 48 hours, revealing EGFR downregulation only upon bsAb treatment.

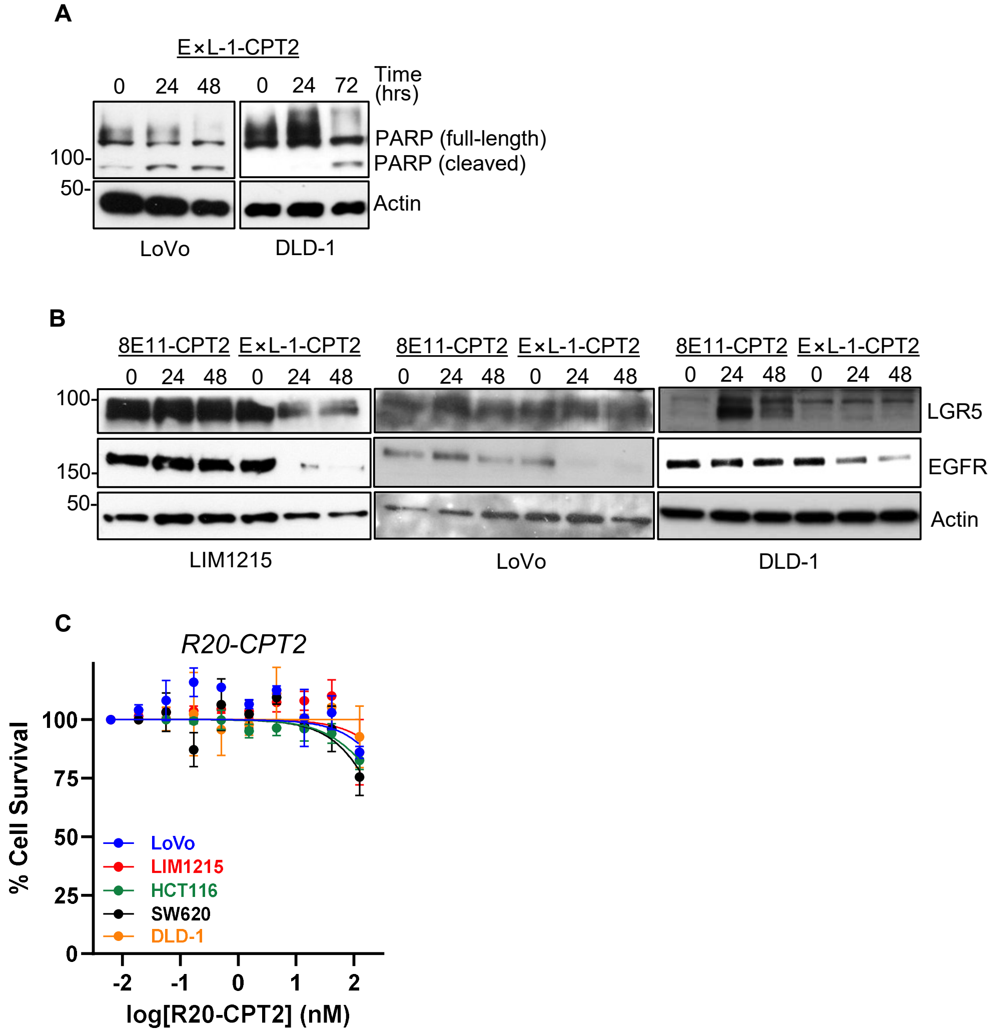


**Figure S3. E×L-1-CPT2 effects on PARP cleavage and receptor expression and R20-CPT2 cytotoxicity.** (**A**) LoVo and DLD-1 cells were treated with 1 µg/ml or 3 µg/ml E×L-1-CPT2, respectively, for the indicated time courses. (**B**) LIM1215, LoVo and DLD-1 cells were treated with 1 µg/ml 8E11-CPT2 or E×L-1-CPT2 for the indicated time courses, revealing EGFR downregulation only upon bsADC treatment. (**C**) R20-CPT2 cytotoxicity in a panel of cancer cell lines after 4 days.

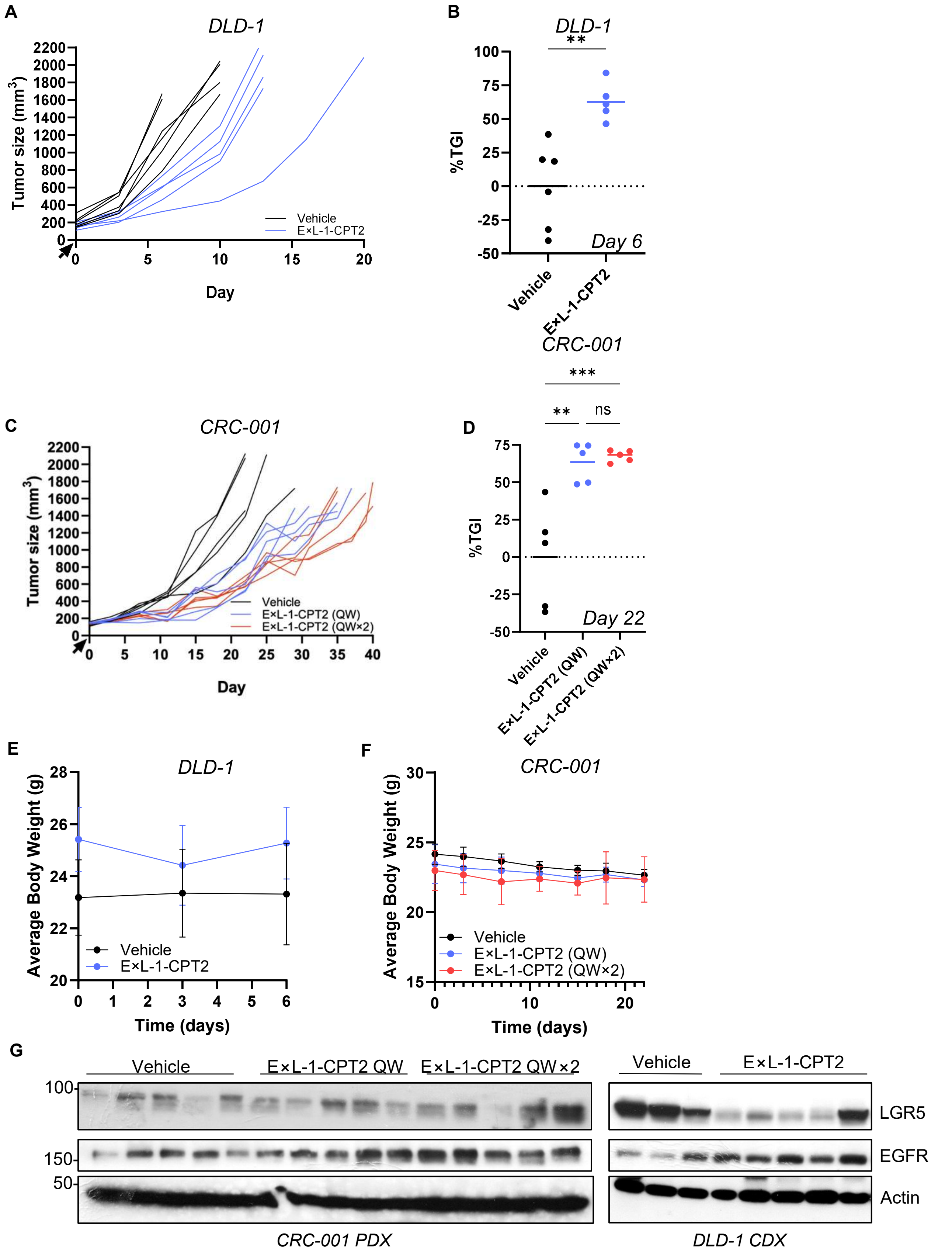


**Figure S4. Individual tumor growth curves, TGI, body weights, and biomarker expression for DLD-1 and CRC-001 PDX studies.** (**A**) Individual tumor curves and (**B**) % TGI from DLD-1 CDX study. (**C**) Individual tumor curves and (**D**) % TGI from CRC-001 PDX study. Average body weights from (**E**) DLD-1 and (**F**) CRC-001 xenograft studies. Data presented as mean +/- SD. (**G**) Western blot of LGR5 and EGFR levels in CRC-001 and DLD-1 xenograft tumors collected at maximal tumor burden. All tumors retained EGFR and/or LGR5 expression.

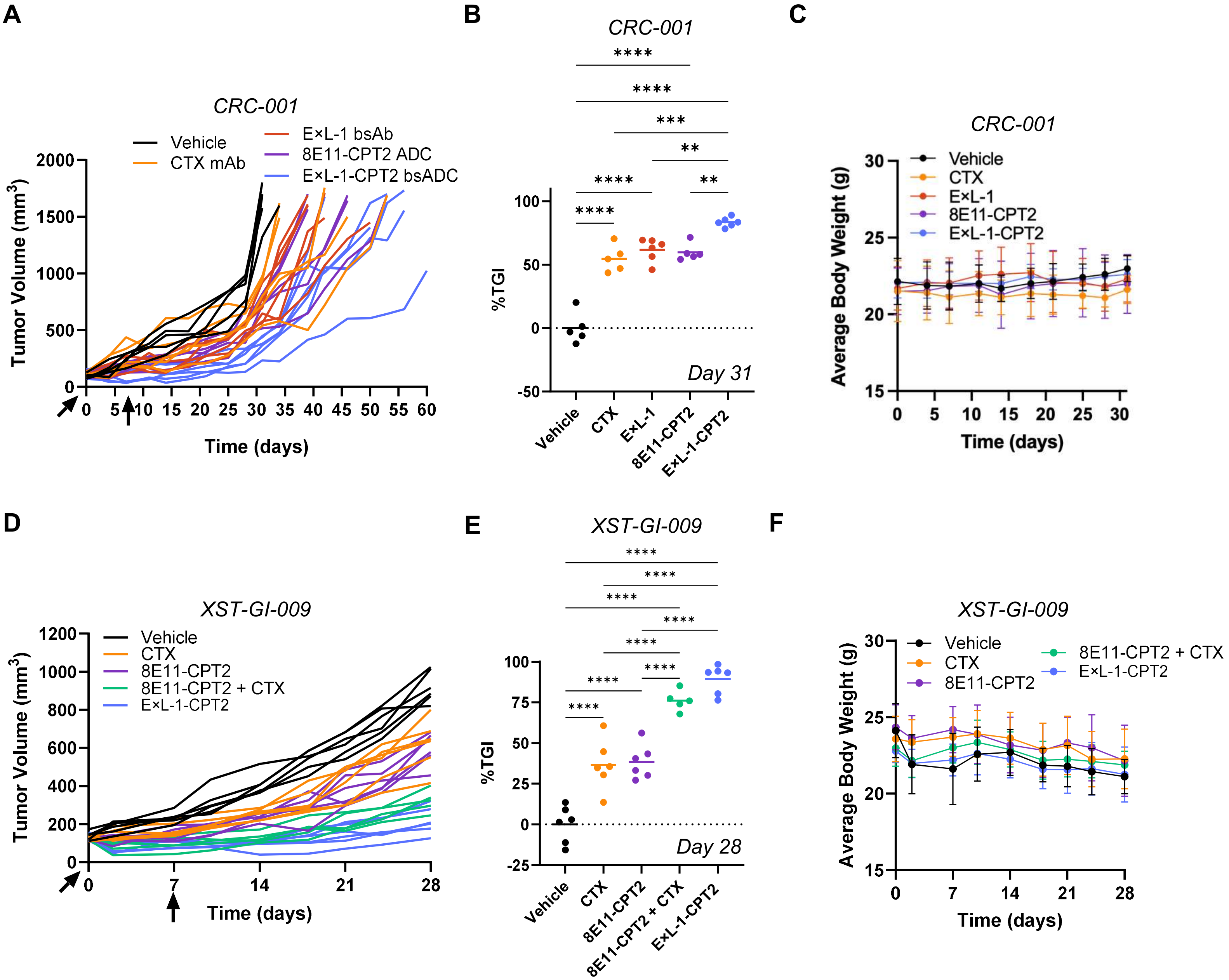


**Figure S5. Individual tumor growth curves, TGI, and body weights for CRC-001 and XST-GI-009 PDX studies.** (**A**) Individual tumor growth curves, (**B**) % TGI, and (**C**) average body weights from CRC-001 PDX study examining E×L-1-CPT2 efficacy against CTX, E×L-1, and 8E11-CPT2. (**D**) Individual tumor growth curves, (**E**) average body weights, and (**F**) % TGI from XST-GI-009 PDX study examining E×L-1-CPT2 efficacy against CTX, 8E11-CPT2, and CTX + 8E11-CPT2 combination.

| | $K_D$ +/- SD (nM) | |
|---|---|---|
| *mAb/bsAb* | *EGFR* | *LGR5* |
| CTX | 0.392 +/- 0.247 | ND |
| 8E11 | ND | 1.290 +/- 0.142 |
| E×L-1 | 5.692 +/- 1.352 | 1.364 +/- 0.873 |
| E×L-2 | 3.976 +/- 1.911 | 2.103 +/- 0.372 |

**Table S1. EGFR and LGR5 mAb and bsAb $K_D$ values.** Average $K_D$ values for CTX, 8E11, E×L-1, and E×L-2. Data presented as mean +/- SD. ND: not determined

| $IC_{50}$ +/- SD (nM) | |
|---|---|
| *Cell line* | *CPT2* |
| HCT116 | 0.79 +/- 0.5826 |
| AGS | 2.829 +/- 0.279 |
| LoVo | 18.66 +/- 11.28 |
| SW948 | 678.7 +/- 407.4 |
| LIM1215 | 27.50 +/- 23.27 |
| DLD-1 | 20.14 +/- 5.101 |
| HT-29 | 111.76 +/- 56.02 |
| SK-N-AS | 1.07 +/- 1.019 |
| SK-N-BE(2) | 12.50 +/- 5.407 |
| SK-CO-1 | 349.4 +/- 308.2 |

**Table S2. CPT2 $IC_{50}$ values.** Average CPT2 payload $IC_{50}$ values for a panel of cancer cell lines. Data presented as mean +/- SD.

| | **$IC_{50}$ +/- SD (nM)** | |
|---|---|---|
| *Cell line* | *E×L-1-CPT2* | *8E11-CPT2* |
| AGS | 0.177 +/- 0.171 | 62.32 +/- 12.28 |
| LIM1215 | 0.125 +/- 0.0714 | 124.75 +/- 24.66 |
| LoVo | 0.205 +/- 0.167 | 53.77 +/- 37.73 |
| DLD-1 | 1.382 +/- 1.706 | 209.81 +/- 32.63 |
| SK-CO-1 | 3.11 +/- 1.00 | 39.14 +/- 8.20 |
| SW948 | 5.79 +/- 2.21 | 235.19 +/- 2.63 |
| HCT116 | 0.406 +/- 0.187 | ND |
| HT-29 | 1.03 +/- 0.388 | ND |
| SW620 | 39.52 +/- 12.37 | 61.62 +/- 5.95 |
| SW620-EGFR | 0.0928 +/- 0.0608 | 34.11 +/- 29.70 |
| SK-N-AS | 0.0271 +/- 0.0185 | 21.15 +/- 12.34 |
| SK-N-BE(2) | 0.0533 +/- 0.0199 | 14.204 +/- 8.98 |

**Table S3. ADC $IC_{50}$ values.** Average E×L-1-CPT2 and 8E11-CPT2 ADC $IC_{50}$ values for a panel of cancer cell lines. Data presented as mean +/- SD. ND: not determined
